# AAGGG repeat expansions trigger RFC1-independent synaptic dysregulation in human CANVAS Neurons

**DOI:** 10.1101/2023.12.13.571345

**Authors:** Connor J. Maltby, Amy Krans, Samantha J. Grudzien, Yomira Palacios, Jessica Muiños, Andrea Suárez, Melissa Asher, Vikram Khurana, Sami J. Barmada, Anke A. Dijkstra, Peter K. Todd

**Author notes:** Corresponding Author. Current address: Peter K. Todd, Department of Neurology, University of Michigan, Ann Arbor, MI, 48109, USA.

## Abstract

Cerebellar ataxia with neuropathy and vestibular areflexia syndrome (CANVAS) is a late onset, recessively inherited neurodegenerative disorder caused by biallelic, non-reference pentameric AAGGG(CCCTT) repeat expansions within the second intron of replication factor complex subunit 1 (*RFC1*). To investigate how these repeats cause disease, we generated CANVAS patient induced pluripotent stem cell (iPSC) derived neurons (iNeurons) and utilized calcium imaging and transcriptomic analysis to define repeat-elicited gain-of-function and loss-of-function contributions to neuronal toxicity. AAGGG repeat expansions do not alter neuronal RFC1 splicing, expression, or DNA repair pathway functions. In reporter assays, AAGGG repeats are translated into pentapeptide repeat proteins that selectively accumulate in CANVAS patient brains. However, neither these proteins nor repeat RNA foci were detected in iNeurons, and overexpression of these repeats in isolation did not induce neuronal toxicity. CANVAS iNeurons exhibit defects in neuronal development and diminished synaptic connectivity that is rescued by CRISPR deletion of a single expanded allele. These phenotypic deficits were not replicated by knockdown of RFC1 in control neurons and were not rescued by ectopic expression of RFC1. These findings support a repeat-dependent but RFC1-independent cause of neuronal dysfunction in CANVAS, with important implications for therapeutic development in this currently untreatable condition.

**Summary:** Human CANVAS neurons exhibit transcriptional and functional synaptic defects that are corrected by heterozygous repeat deletion but are independent of the gene within which they reside—RFC1.

## Introduction

Cerebellar ataxia with neuropathy and vestibular areflexia syndrome (CANVAS) is a recessively inherited, progressive, and debilitating disorder characterized by vestibular, cerebellar, and somatosensory impairments^1,2^. CANVAS typically presents in late-middle-age, with most patients exhibiting progressive motor imbalance, oscillopsia, dysphagia, dysarthria, and peripheral sensory neuropathy. Post-mortem analyses of CANVAS patients indicate cerebellar and basal ganglia atrophy with diffuse Purkinje cell loss, as well as peripheral sensory neuronopathy including the vestibular, facial, and trigeminal nerves^1,3–7^ (Figure 1A).

**Figure 1.**
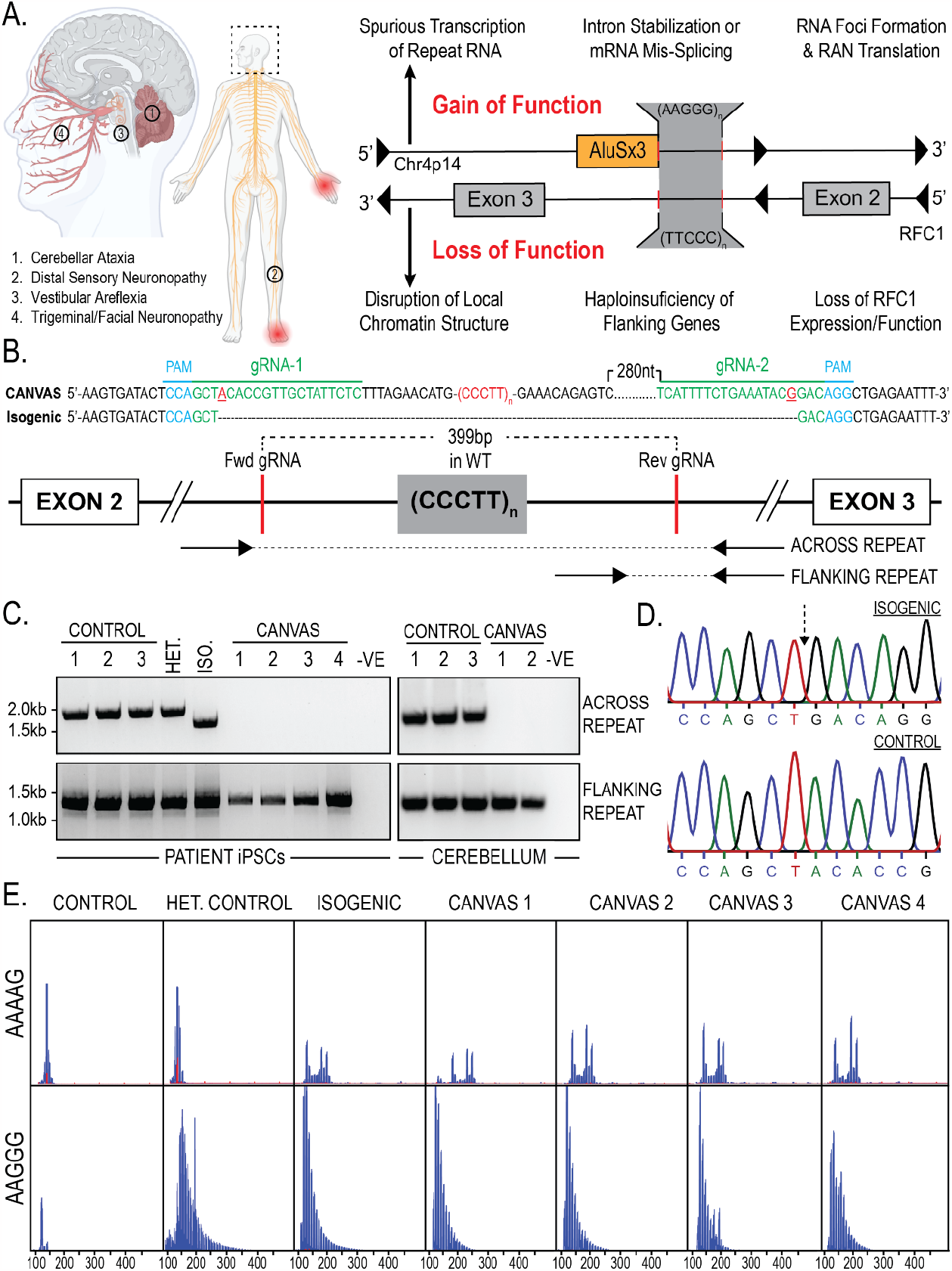
Repeat characterization and heterozygous correction of CANVAS patient derived iPSC lines. **A**) Schematic of brain, CNS, and PNS regions affected in CANVAS (left), and potential mechanisms of repeat toxicity in CANVAS (right). **B**) Repeat architecture of the expanded locus and CRISPR gRNA design to remove the AAGGG/CCCTT repeat expansion by NHEJ. **C**) Endpoint PCR of gDNA extracted from CANVAS patient and control derived iPSC lines and CANVAS and control cerebellum tissue utilizing the primer pairs outlines in (1B) to screen for the presence of WT repeat, mutant repeat expansion, or deletion of expanded repeat. **D**) Chromatogram of Sanger Sequencing identifying AAGGG/CCCTT allele deletion in heterozygous isogenic line indicating the expected NHEJ join point compared to control iPSC line. **E**) Repeat Primed PCR of the expanded locus in CANVAS patient and control derived iPSC lines utilizing anti AAAAG and AAGGG probes.

CANVAS results from a biallelic, non-reference, pentameric CCCTT(AAGGG) repeat expansion in the second intron of replication factor complex subunit 1 (*RFC1*) and this same expansion also causes late-onset idiopathic ataxia and sensory neuropathy in isolation^8–10^. Normally, this locus harbors a short (AAAAG)_∼11_ repeat, and individuals harboring a heterozygous (AAGGG)_exp_ allele with the wild type (AAAAG)_∼11_ allele do not develop CANVAS spectrum phenotypes, consistent with a recessive pattern of inheritence^8,9^. Since the initial description of AAGGG expansions at this locus, several further pathogenic repeat expansion conformations have been described: ACAGG expansions have been identified in CANVAS patients of Oceania and East Asian descent^11^, and more recently the 100,000 genomes project database has identified AAGGC, AGGGC, and AGAGG expansions, either in the homozygous or compound heterozygous state with the AAGGG expansion^12^. Furthermore, AAAGG repeat expansions previously reported to be non-pathogenic in the homozygous state and compound heterozygous state with AAGGG expansions have since been found to be pathogenic when present at significantly expanded states^12^. *RFC1*-associated repeat expansions are common, with a minor allelic frequency of ∼0.7-6.8% and an expected homozygous pathogenic repeat population frequency of 1 in 625 individuals worldwide—making it one of the most common causes of inherited ataxia and sensory neuropathy^8,13,14^

Despite clinical interest in CANVAS, the mechanisms by which this repeat expansion causes pathogenesis and neuronal death are unknown. Initial studies suggested that *RFC1* mRNA and protein expression are unaltered in the context of the repeat expansion, which is inconsistent with the classical mechanism by which recessively inherited repeat expansions elicit a loss of function for the genes in which they reside^8,10,15,16^. However, identification of rare compound heterozygotic CANVAS patients harboring *RFC1* truncating and splicing loss-of-function mutations with single allele repeat-expansions suggests a role for RFC1 function in CANVAS^17–19^ . Alternatively, the repeats could potentially elicit toxicity through dose-dependent gain of function mechanisms (such as repeat associated non-AUG initiated (RAN) translation or repeat RNA-protein complex formation) that only manifest in homozygosity or in combination with RFC1 haploinsufficiency^20,21^ (Figure 1A). Limited postmortem and cell-based analyses performed to date have not demonstrated classic pathological hallmarks seen in other repeat expansion disorders, such as repeat RNA foci or ubiquitinated inclusions^8^.

Understanding the pathogenic nature of this repeat expansion is further complicated by its manifestation within two distinct genetic elements on opposing strands. The AAGGG repeat sits at the 3’ end of an AluSx3 transposable element on the sense strand, while the complementary CCCTT repeat is embedded deep within the large intron 2 region of *RFC1*, potentially allowing for two distinct and context-dependent mechanisms of toxicity. Alu elements are ancient retro-transposition artifacts that make up roughly 10% of the human genome (∼1 million copies)^22^ and serve as reservoirs of potential regulatory functions that have actively driven primate evolution, with evidence to suggest roles for intronic Alu elements in mRNA splicing^23,24^ (constitutive and alternative), RNA editing^25^, and protein translation^26^. Moreover, the AAGGG repeat sequence found on the opposing strand to *RFC1* at this AluSx3 3’ end is predicted to form G-quadruplex secondary structures implicated in repeat expansion disease pathogenesis in both transcribed RNA and DNA during transcription and DNA replication fork formation^27^.

Given the paucity of empiric data on these repeats and the multiple potential mechanisms underpinning CANVAS, we investigated how these repeats cause disease, including independent evaluation of previously tested ideas and direct measures of the ability of these repeats to elicit neurotoxicity. We generated multiple CANVAS patient-derived and CRISPR-corrected isogenic induced pluripotent stem cell (iPSC) lines and differentiated them into neurons.

Through functional neuronal assays, transcriptomic analyses, and *in vitro* toxicity investigations, we identify deficits in key neuronal cellular pathways in CANVAS neurons that are rescued upon CRISPR correction of a single expanded repeat allele. In contrast, loss of RFC1 expression fails to recapitulate these cellular and molecular phenotypes and reprovision of RFC1 into CANVAS iNeurons is insufficient to reverse pathologic cascades. Together, these studies support a repeat-dependent mechanism of toxicity that operates outside of the canonical functions of RFC1.

## Results

### CANVAS patient-derived and CRISPR-corrected heterozygous isogenic iPSC lines

Homozygous non-reference repeat expansions in *RFC1* could potentially elicit neurodegeneration through multiple gain-of-function or loss-of-function mechanisms (Figure 1A). To assess these possibilities systematically, we generated a series of CANVAS patient derived-derived iPSC lines from 4 patients and controls, as well as one family member with a heterozygous *RFC1* expansion (Supplementary Figure 1A-B). As an additional experimental control, we generated an isogenic line through deletion of the expanded allele in patient one iPSC line through CRISPR-Cas9 genome editing (Figure 1B-D). Successful deletion of a single expanded allele was achieved in CANVAS patient 1 iPSCs (Figure 1C), and the specificity of this deletion was confirmed by Sanger sequencing of PCR products (Figure 1D). However, we were unable to generate biallelic homozygous deletions of this AluSx3 element and expanded repeat despite multiple rounds of editing and re-editing in both CANVAS and control cell lines, consistent with prior reports^28^. As such, the heterozygous deletion line was taken forward for experimental analysis. Cell lines were confirmed to possess AAGGG/CCCTT repeat expansion alleles by repeat-primed PCR (Figure 1E) in combination with the results obtained by screening PCR (Figure 1C).

### Translated AAGGG repeat products are detected in CANVAS patient brains

To investigate the potential of repeat-dependent gain-of-function mechanisms in CANVAS pathogenesis (Figure 1A), we generated repeat-containing reporter constructs encompassing various repeat motifs within intronic sequence contexts (Figure 2A). Wild-type AAAAG/CTTTT and mutant CANVAS AAGGG/CCCTT repeat expansions were generated by recursive directional ligation^29^ and were ligated into plasmid backbones containing 150bp of upstream intronic sequence deriving from the genomic strand within which the repeat motif would be found; AAAAG/AAGGG repeats for the sense strand AluSx3 containing sequence and CTTTT/CCCTT repeats for the antisense *RFC1* intron 2 containing sequence.

**Figure 2.**
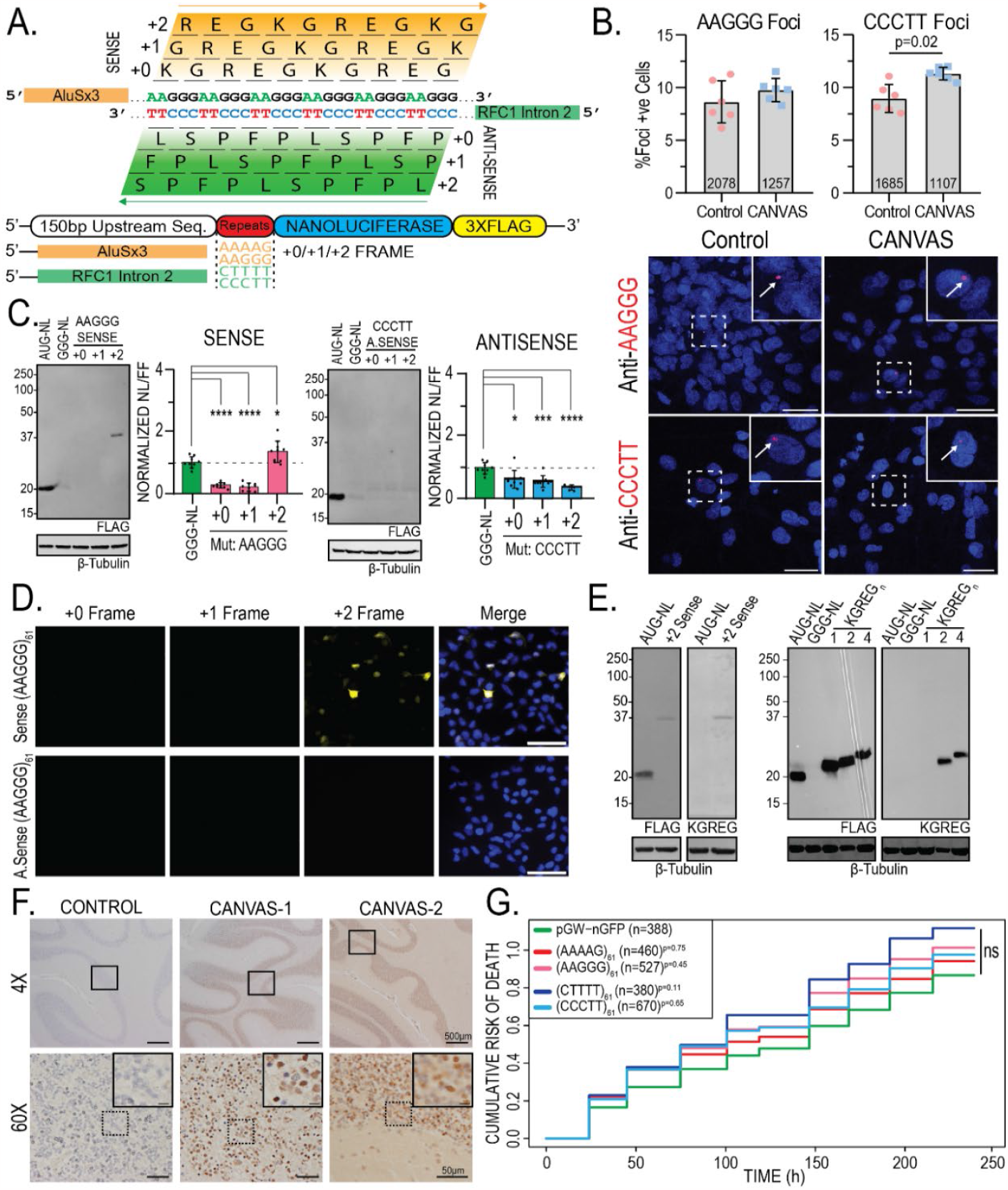
Translated AAGGG repeat products are detected in CANVAS patient brains. **A)** Schematic of reading frames and peptide products from the sense and antisense strands of the repeat expansion locus. **B)** Confocal Images of representative CANVAS patient and control neurons after RNA HCR with anti-AAGGG or anti-CCCTT fluorescent probes, scale = 10 μm, and quantification of foci positive neurons for control (n=3) and CANVAS (n=3) patient neurons with total n-numbers of neuronal cells analyzed indicated. AAGGG (F(5, 12) = 3.619, P=0.074), CCCTT (F(5, 12) = 8.293, P=0.011). n = 2 biological replicates from 3 independent patient derived cell lines. Data were analyzed by one-way ANOVA with Sidak’s post-hoc multiple comparison tests. **C**) Expression analysis of lysates from HEK293 cells transfected with control plasmid or plasmids encoding intronic sense or antisense AAGGG/CCCTT repeat reporters in the +0/+1/+2 reading frames (left) and Nano-luciferase expression assay quantification (right). N = 3 technical replicates from a total of 7 biological replicates. Data were analyzed by one-way ANOVA with Sidak’s post-hoc multiple comparison tests. **D**) ICC of HEK293 cells transfected with plasmids encoding intronic sense or antisense AAGGG/CCCTT repeat reporters with C-terminal triple-tags in the +0/+1/+2 reading frames. **E**) Expression analysis of lysates from HEK293 cells transfected with control plasmid +2 Sense (AAGGG)_61_ plasmid using anti-FLAG M2 (1:1000) and anti-KGREG (1:100) antibodies (left). **F**) IHC of control (n=1) and RFC1 expansion CANVAS (n=2) patient post-mortem cerebellar vermis tissue stained with sense anti-KGREG antibody (1:100, acid AR). Scale = 500 μm (4x), 50 μm (60x) and 20 μm (inset). Full IHC results of 3x control tissues are shown in supplementary figure 2C. **G**) Cumulative Hazard Plot for primary rat cortical neurons transfected with CANVAS intronic expression plasmids containing 61 repeats of the indicated type over a 10-day period, n = 8 technical replicates, 3 biological replicates, n numbers of cells shown per condition. Error = SD.

We first assessed whether expression of these repeat containing constructs elicit RNA foci formation by using repeat-targeting fluorescent probes and RNA HCR-FISH (Supplementary Figure 2A). In HEK293 cells, sense (AAGGG) repeat-directed probes detected nuclear and cytoplasmic RNA foci that co-localized with anti-nanoluciferase probes in transfected cells and generated patterns of expression that were different from nanoluciferase expression alone. Most of the nuclear and all the cytoplasmic signal was ablated by RNase treatment (Supplementary Figure 2A). A similar pattern was seen for antisense (CCCTT) repeat-directed probes, suggesting that both the sense and antisense pentanucleotide repeats can form RNA condensates within cells. To assess if such foci are detectable in patient cells, we performed RNA HCR-FISH with these same probes in control (n=3) and CANVAS (n=3) patient iPSC-derived neurons (Figure 2B, Supplementary Figure 2B-C). No significant difference was observed between CANVAS and control neurons for sense AAGGG repeat-RNA foci detection (F(5, 12) = 3.619, P=0.074). In contrast, antisense CCCTT repeat-RNA foci were detected in CANVAS neurons at greater rates compared to control cells (F(5, 12) = 8.293, P=0.011). However, the overall abundance of foci was low, and the specificity was imperfect, with 8.94% of control neurons and 11.3% of CANVAS neurons showing antisense CCCTT RNA foci in blinded assays (Figure 2B, Supplementary Figure 2D).

The native sequences surrounding the repeats are predicted to contain both AUG and non-AUG near-cognate codons upstream of the repeats without any intervening stop codons—potentially placing the repeats within open reading frames. To assess whether these repeats might be translated in patients, we generated intronic sense and antisense reporter constructs with Nanoluciferase-3xFlag (NL-3F) C-terminal tags in all 3 potential reading frames.

Protein expression and nanoluciferase activity analysis in transfected HEK293 cells showed selective translation of repeat-derived peptides within the sense (AAGGG) strand +2 reading frame (Figure 2C, left). In contrast, no products were detected from the antisense *RFC1* intron in all 3 reading frames (Figure 2C, right), despite the presence of an AUG codon in the +0 reading frame. This was also observed by ICC in HEK293 cells, whereby only cells transfected with the +2 sense AAGGG reporter showed translation of any repeat-containing peptides (Figure 2D).

If the AAGGG repeat were to be translated into a protein, it would generate the same pentapeptide repeat protein, polyKGREG, in all 3 potential reading frames. We therefore generated antibodies against a polyKGREG epitope. This antibody showed a high degree of specificity for two or more KGREG repeats, which are otherwise not predicted to occur in the human proteome, in transfected HEK293 cells by immunoblot staining (Supplementary Figure 3A) and strong colocalization with FLAG antibodies by immunocytochemistry (Supplementary Figure 3B). Using this antibody, we confirmed that the +2 sense AAGGG reporter product contained KGREG peptides (Figure 2E). To assess whether sense-strand derived KGREG peptide products accumulate in CANVAS patient brains, we performed immunohistochemistry on control (n=3) and CANVAS (n=2) post-mortem brain samples obtained from the University of Michigan Brain Bank, Harvard University Brain Bank, and the University of Amsterdam Brain Bank (Figure 2F, Supplementary Figure 3C). KGREG staining was selectively detected within cerebellar granule cells in CANVAS patients, with no signal detected in controls. While CANVAS patient brains showed marked Purkinje cell loss, no KGREG staining was observed in the few Purkinje cells that remained (Figure 2F, Supplementary Figure 3C). These data suggest that pentapeptide KGREG repeat proteins may be produced from AAGGG repeats in CANVAS patients in a cell type-specific manner.

Expression of either CGG or GGGGCC repeats in neurons is sufficient to elicit toxicity, which can serve as a proxy for gain-of-function associated neurodegeneration^30–34^. We therefore tested whether ectopic AAGGG or CCCTT repeat expression might elicit neurotoxicity. Using automated longitudinal fluorescence microscopy^35,36^, expression of either the intronic WT or mutant sense or antisense reporter constructs in primary rat cortical neurons showed no toxicity over a 10-day period compared to expression of GFP alone (AAAAG_61_: P=0.75, AAGGG_61_: P=0.45, CTTTT_61_: P=0.11, CCCTT_61_: P=0.65) (Figure 2G). These data suggest that while AAGGG and CCCTT repeats are capable of eliciting RNA foci and being translated into pentapeptide repeat proteins, they do not exhibit intrinsic toxicity and do not accumulate into pathologic aggregates in patient tissues.

### Splicing of RFC1 is normal in multiple CANVAS patient derived cell types

Intronic repeat expansions can interfere with pre-mRNA splicing by inducing intron retention within the mature transcript ^37,38^, stabilizing circular intronic lariat species^39^, or inducing differential exon usage during splicing^40–42^. Intronic Alu elements also influence alternative splicing through inclusion of Alu-containing exons, with 85% of Alu containing exons deriving from antisense Alu elements^43,44^. To assess whether AAGGG/CCCTT repeat expansions induced intron retention, we performed end-point RT-PCR with primers that amplify the exon2-exon3 junction or the exon2-intron2 junction (Figure 3A). There were no differences between cases and controls in terms of intron retention or aberrant intron 2 splicing of *RFC1* in patient fibroblasts, iPSC-derived neurons or cortical and cerebellar regions of post-mortem brain (Figure 3A).

**Figure 3.**
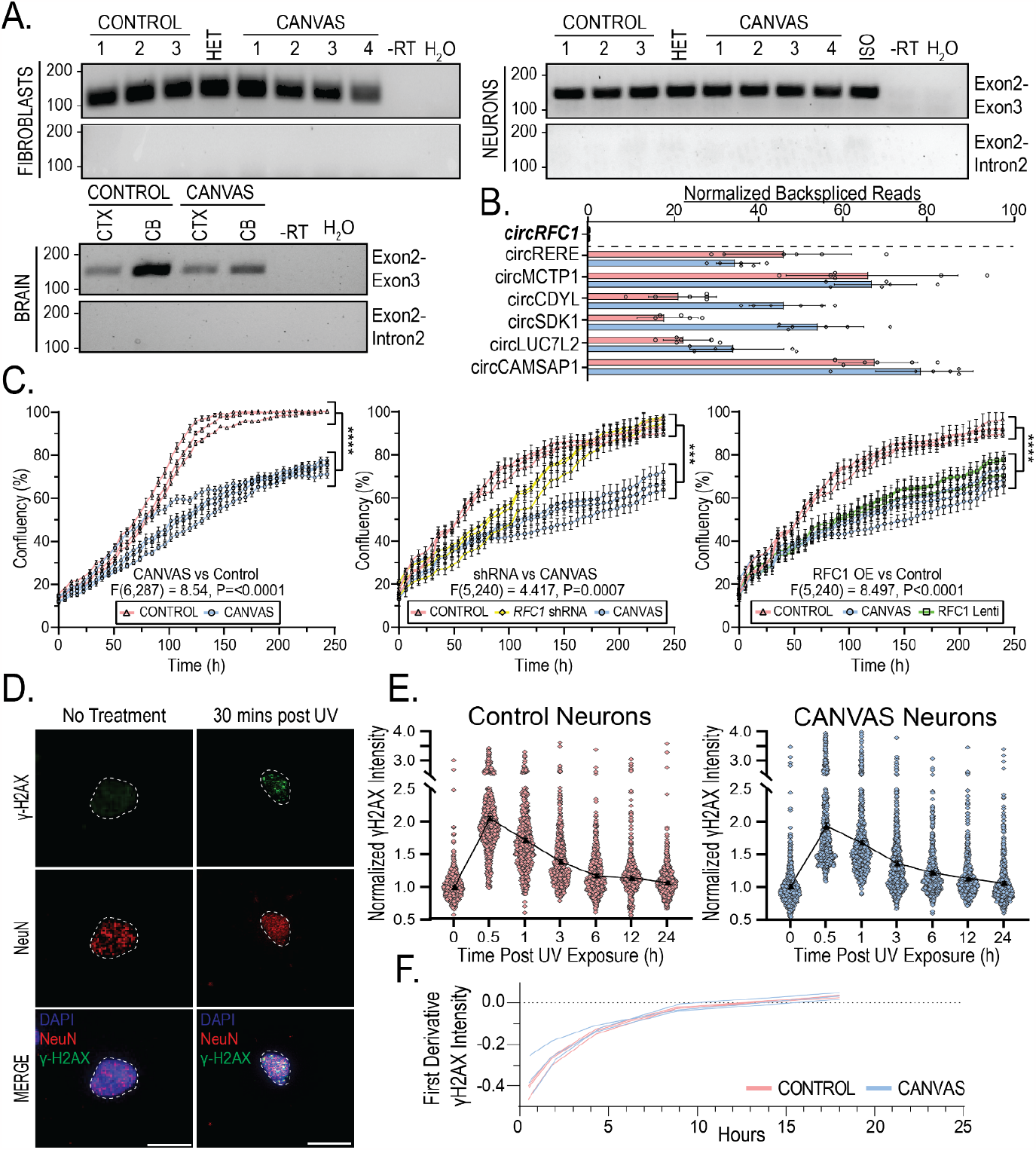
Canonical functions and expression of RFC1 are normal in CANVAS patient-derived cells. **A)**Endpoint RT-PCR utilizing primer sets spanning RFC1 exon2-exon3 or exon2-intron 2 in CANVAS fibroblasts (Top, left), iPSC-derived neurons (Top, right), and CANVAS post-mortem brain (Bottom, left). **B**) Quantification of normalized circular backspliced read counts for *RFC1* and other known circRNA species in CANVAS patient iPSC-derived neurons by paired-end RNASeq analysis. **C**) 10-day time-course analysis of the rate of cellular division and proliferation in CANVAS (n=4) and control (n=3) fibroblast lines (**left**, F(6,287) = 8.54, P<0.0001), control fibroblast lines (n=3) mock-treated or treated with RFC1 shRNA lentivirus (**center**, F(5,240) = 1.314, P=0.131), and CANVAS fibroblast lines (n=3) mock-treated or treated with RFC1 overexpression lentivirus (**right**, F(5,240) = 2.358, P=0.245). N = 3 biological replicates from 3-4 independent patient cell lines. Data were analyzed by one-way ANOVA with Sidak’s post-hoc multiple comparison tests. **D-E**) Analysis of recovery after discrete UV exposure and DNA damage in CANVAS patient iPSC-derived neurons. **D**) Representative images of γ-H2AX staining of iPSC-derived neurons pre- and post-60 mJ/cm^2^ UV exposure (Scale = 10μm). **E**) Quantification of mean γ-H2AX staining in CANVAS patient (n=3) and control (n=3) iPSC-derived NeuN+ neuronal nuclei over a 24h period after 60 mJ/cm^2^ UV exposure (n=16,569 NeuN+ nuclei total). Data were analyzed by one-way ANOVA with post-hoc multiple comparison tests. **F**) First derivative DNA damage recovery rate curves for CANVAS (n=3) and control (n=3) patient iPSC-derived neurons. Error = SD.

To investigate whether AAGGG/CCCTT repeat expansions trigger *RFC1* intron 2 stabilization as a circular lariat species, as may occur in *C9orf72* FTD/ALS^39^, we utilized whole-transcriptome RNA short read sequencing to map *RFC1* mRNA splicing isoforms and identify the presence of intronic back-spliced reads indicative of circular mRNA species (circRNAs) arising from circular intronic lariats^45^. Total rRNA-depleted RNA was extracted from 10-week-aged CANVAS patient and control iPSC-derived neurons and paired-end reads were processed and analyzed using packages described in methods. A known list of circRNA species^46–50^ were detected at comparable levels in CANVAS and control samples (Figure 3B), however, no backspliced reads were identified to map to *RFC1* intron 2 region or across the *RFC1* transcript, indicating a lack of detectable circular RNA species deriving from the *RFC1* locus (Figure 3B).

To assess whether AAGGG/CCCTT repeat expansions affect *RFC1* mRNA isoforms, we used DEXSeq^51^ to assess the normalized differential exon usage of mature spliced *RFC1* transcripts in CANVAS patient and control iPSC-derived neurons (Supplementary Figure 4A). No changes in *RFC1* exon usage were observed within the N-terminal regions flanking the repeat expansion, or across the entire *RFC1* transcript. Similarly, Sashimi plots of *RFC1* splicing demonstrate similar splicing and alternative exon usage in CANVAS patient iPSC-derived neurons compared to controls without evidence for alternative non-canonical exon usage for *RFC1* in normal or disease states (Supplementary Figure 4B).

### Canonical functions and expression of RFC1 are normal in CANVAS patient-derived cells

Prior studies in CANVAS patient fibroblasts, lymphoblasts, frontal cortex and cerebellar vermis of CANVAS patients demonstrated no differences in *RFC1* mRNA when compared to controls^8^. Consistent with prior studies, we observed no changes in *RFC1* steady-state mRNA abundance or RFC1 protein expression between control (n=3), CANVAS (n=4), and heterozygous carrier patient fibroblasts (n=1) (Supplementary Figure 5A). Similarly, we observed no changes in *RFC1* mRNA or RFC1 protein levels between 6-week-old control (n=3) and CANVAS (n=3) patient iPSC-derived neurons (Supplementary Figure 5B). Analysis in limited post-mortem frontal cortex and cerebellar brain samples showed no change in *RFC1* mRNA, with a modest but significant increase in RFC1 protein (Supplementary Figure 5C).

RFC1 plays critical roles in both DNA replication and DNA damage repair pathways as a subunit of the DNA clamp-loader complex^52–54^. To assess whether AAGGG/CCCTT repeat expansions might interfere with RFC1’s DNA replication functions, we first investigated the rate of cellular proliferation in age- and passage-matched CANVAS (n=4) and control (n=3) patient-derived fibroblasts (Figure 3C, left). We observed a highly significant decrease in cellular proliferation rate for CANVAS fibroblasts when compared to controls (F(6,287) = 8.54, P<0.0001). To determine if this CANVAS fibroblast proliferation phenotype was dependent on RFC1 expression or function, control-derived fibroblasts (n=3/group) were transduced with lentiviral particles encoding shRNAs targeting *RFC1* exon 4 and exon 15, or a control shRNA, and were assessed for proliferation rate over a 10-day period as before (Figure 3C, Center). *RFC1* knockdown reduced RFC1 protein to nearly undetectable levels (Supplementary Figure 7). RFC1 knockdown in control fibroblasts resulted in an initial lag in cellular proliferation compared to control shRNA-treated fibroblasts (120h, P= 0.001) which recovered to control levels after 200h (Figure 3C, Center, F(5,240) = 1.314, P=0.131), while untreated CANVAS patient fibroblasts continued to show a slower proliferation rate throughout the analysis (F(5,240) = 4.417, P=0.0007). To assess if re-provision of full-length wild-type RFC1 was sufficient to overcome the CANVAS phenotype, we overexpressed full-length RFC1 in CANVAS patient-derived fibroblasts (Figure 3C, Right). CANVAS patient-derived fibroblasts overexpressing RFC1 retained a markedly slowed proliferation rate compared to control fibroblasts (F(5,240) = 8.497, P<0.0001), with no significant improvement over the proliferation rate of CANVAS patient-derived fibroblasts treated with a control lentivirus (F(5,240) = 2.358, P=0.245). These results suggest that AAGGG/CCCTT repeat expansions affect cell proliferation independent of RFC1 protein levels.

CANVAS is predominantly a neuronopathy, and as neurons are post-mitotic, we investigated whether DNA damage repair was dysfunctional in CANVAS patient iPSC-derived neurons. To do this, we analyzed expression of the DNA Damage marker γ-H2AX in 8-week-old control and CANVAS iPSC-derived neurons where basal levels of DNA damage accumulation was comparable between control and CANVAS iPSC-derived neurons (Supplementary Figure 6A). To test for an altered response to DNA damage, CANVAS patient and control iPSC-derived neurons were exposed to UV irradiation (Figure 3D-E). To test for an altered response to DNA damage, CANVAS patient and control iPSC-derived neurons were exposed to UV irradiation (Figure 3D-E). As expected, γ-H2AX levels increased after exposure to 0-120 mJ/cm^2^ UV irradiation in control iPSC-derived neurons (Supplementary Figure 6B). Control (n=3) and CANVAS patient (n=3) iPSC-derived neurons showed a consistent ∼2-fold increase in γ-H2AX reactivity within 30 mins of 60 mJ/cm^2^ UV irradiation, followed by a slow but consistent decline in γ-H2AX reactivity in all experimental conditions, with return to baseline γ-H2AX reactivity within 6-12h post-irradiation. First-derivative analysis of normalized γ-H2AX indicated no differences in the rate of γ-H2AX recovery after UV induction between CANVAS and control neurons (Figure 3E-F, Supplementary Figure 6C-D), suggesting that DNA damage repair is not affected in CANVAS neurons.

### Synaptic genes are downregulated in CANVAS patient-derived neurons

To assess for differences between CANVAS and control iPSC-derived neurons that might provide some insights into disease pathogenesis, we next conducted paired-end sequencing of total RNA extracted from 10-week-old CANVAS patient (n=6) and control (n=6) iPSC-derived neurons to identify potential dysregulated pathways or genes in CANVAS. An average of ∼20-30M reads were obtained per sample. After trimming, genome alignment, and filtering for low transcript reads (see methods), we observed that 5.9% (1313) of detected genes were upregulated and 11.7% (2630) were downregulated in CANVAS patient iPSC-derived neurons compared to control neurons (Figure 4A). *RFC1* transcripts showed a modest but significant increase in CANVAS iNeurons that was below the prespecified fold-change threshold implemented (Figure 4A), with no RFC1-associated GO-terms identified within overrepresented cellular pathways of these dysregulated transcripts in CANVAS (Figure 4B). Principal component analysis (PCA) clustering of neuronal samples based on gene expression patterns demonstrated discrete and distinct clustering of CANVAS neuronal samples as separate from controls (Figure 4C) with 96% of observable variance seen across principal component 1 (PC1), indicating high variability between control and CANVAS gene expression profiles and a dominant source of variation between conditions. Heatmap analysis of the top dysregulated transcripts detected in CANVAS patient iPSC-derived neurons show a strong preference for reduced expression of transcripts (Figure 4D). Gene ontology (GO) analysis indicated a significant overrepresentation of neuronal signaling processes including synaptic signaling and processes that regulate synaptic signaling within the transcripts downregulated in CANVAS vs control (Figure 4B, Supplementary Figure 8), with many of these transcripts localized to neuronal processes and involved in signaling, channel, or transporter activity. Normalized gene counts for select dysregulated synaptic genes indicates that this reduction in synaptic gene expression is highly significant and encompasses both pre- and post-synaptic genes, as well as genes involved in synaptic organization and signal transduction (Figure 4E). A full analysis of dysregulated pathways is shown in Supplementary Figure 8.

**Figure 4.**
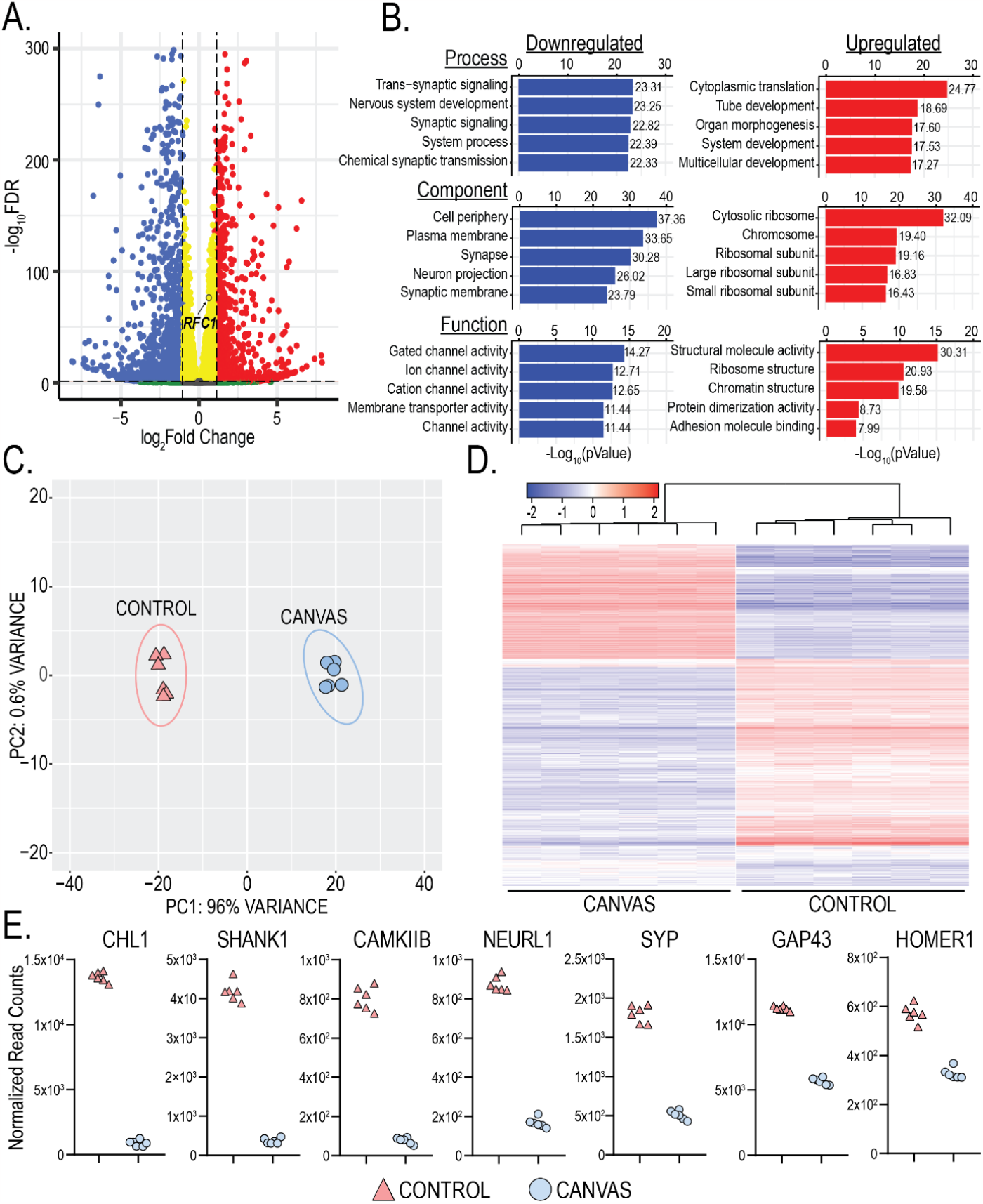
Synaptic genes are downregulated in CANVAS neurons. **A)**Volcano plot of differentially expressed genes in CANVS patient vs control iPSC-derived neurons, blue = significantly downregulated, red = significantly upregulated, RFC1 labelled. **B**) Gene Ontology (GO) pathway analysis of the top 5 up/downregulated Biological Process, Cellular Component, and Molecular Function in CANVAS patient vs control iPSC-derived neurons. N = 6 biological replicates from 3 individual CANVAS and control patients. **C**) Principal Component Analysis (PCA) of CANVAS (n=6) vs control (n=6) patient iPSC-derived neurons. **D**) Heatmap of normalized expression for the top 1000 genes differentially expressed in CANVAS patient vs control iPSC-derived neurons. **E**) Normalized gene counts for the top 7 downregulated synaptic-associated genes in CANVAS patient vs control iPSC-derived neurons.

Protein expression analyses of select synaptic genes significantly downregulated in transcriptomic analyses in Figure 4E found similar downregulation at the protein level. Synaptophysin, CHL1, GAP43, and CAMKIIB all showed highly significant reductions in expression at both the transcript and protein level in CANVAS patient iPSC-derived neurons compared to control (Figure 4E, Figure 5A-D).

**Figure 5.**
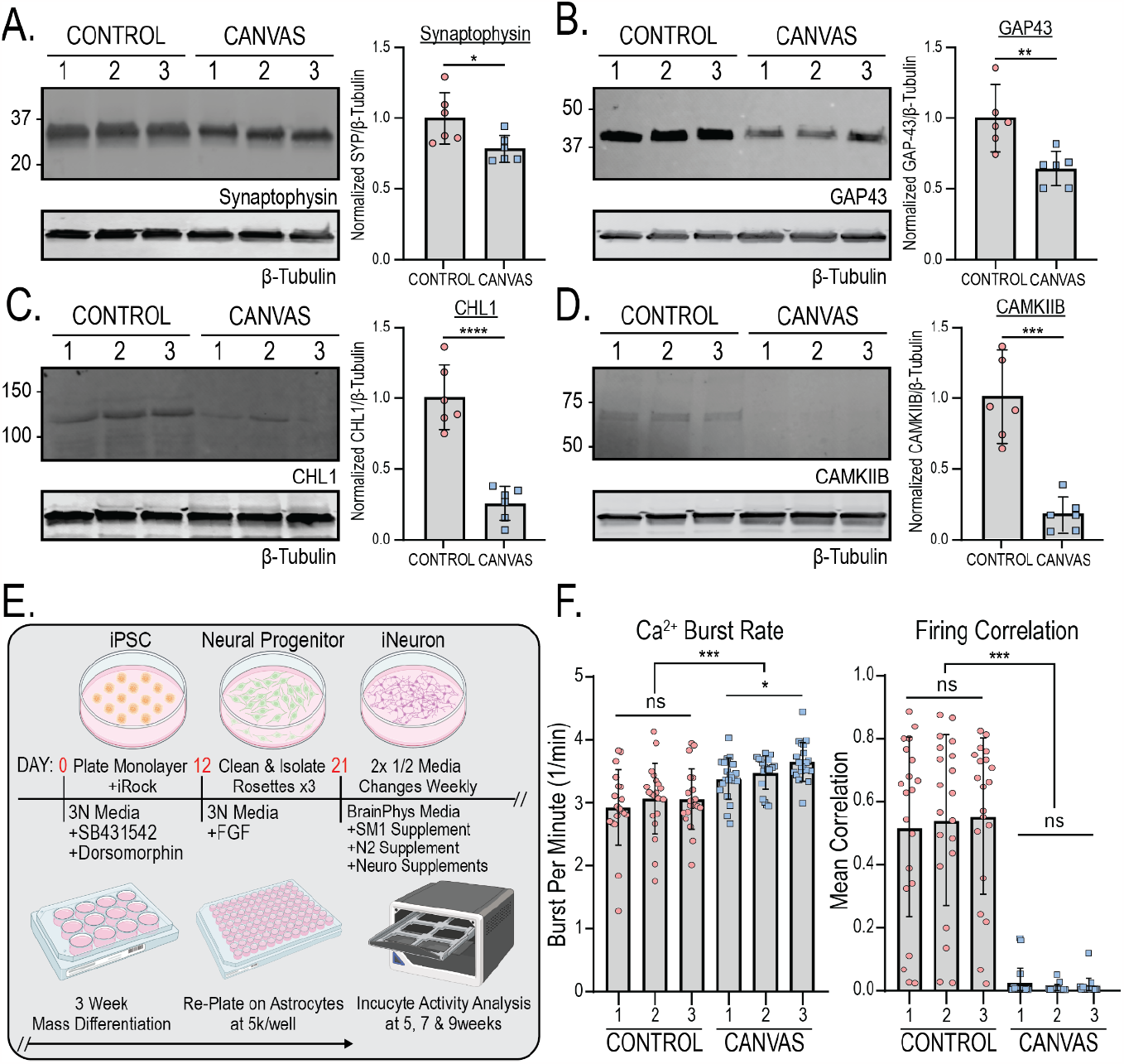
CANVAS patient derived neurons exhibit synaptic dysfunction and reduced connectivity. **A-D**) Protein expression (left) and normalized quantification (right) of selected synaptic genes identified as downregulated in CANVAS patient iPSC-derived neurons by transcriptomic analysis - (**A**) Synaptophysin, (**B**) GAP43, (**C**) CHL1, (**D**) CAMKIIB. N = 2 biological replicates from 3 independent patient derived cell lines. Data were analyzed by one-way ANOVA with Sidak’s post-hoc multiple comparison tests. **E**) Schematic outlining experimental workflow for generating patient iPSC-derived neurons for calcium imaging analysis. **F**) Analysis of Ca^2+^ imaging metrics for control (n=3) and CANVAS (n=3) patient iPSC-derived neurons at 9-weeks post-differentiation. Burst Rate (F(5, 114) = 8.268, P<0.0001), Firing Correlation (F(5, 114) = 45.62, P<0.0001). Each data point represents the mean of ∼1000-3000 active cells per well (Supplementary Figure 5). Data were analyzed by one-way ANOVA with Sidak’s post-hoc multiple comparison tests. Error = SD.

### Spontaneous synaptic activity is impaired in CANVAS patient-derived neurons

Given the dysregulation of synapse-associated transcripts and proteins we observed in CANVAS patient iPSC-derived neurons, we investigated if these alterations in gene expression correlated with observable phenotypic differences in synaptic activity. To accomplish this, we measured spontaneous synaptic activity in control (n=3) and CANVAS (n=3) glutamatergic forebrain neurons with the Incucyte® Neuroburst Orange fluorescent calcium indicator and Incucyte® S3 automated imaging system. Patient-derived neural progenitor cells were differentiated and re-plated at uniform density of 5,000 cells per well on an astrocyte feeder layer within PLO/Laminin coated 96-well plates described in methods (Figure 5E). Neurons were analyzed for active cell number, basal calcium intensity, burst rate, burst strength, burst duration, and network correlation between 3-11 weeks post-differentiation (Figure 5F, Supplementary Figure 9A), and wells with at least 500 active cells were included in the analysis. Both control and CANVAS iPSC-derived neurons consistently had between 1000-2000 active cells per well with minor but statistically significant reductions observed for basal calcium intensity, burst duration, and burst strength in CANVAS neurons (Supplementary Figure 9A). Notably, while control iPSC-derived neurons formed synchronized networks with 50-70% network correlation by 5 weeks post-differentiation (Figures 5F and 6E), CANVAS patient iPSC-derived neurons remained devoid of detectable synchronous firing even 7-11 weeks post-differentiation, with single cells showing independent activity at a 17% increase in firing rate relative to control (Figure 5F, Supplementary Movies).

**Figure 6.**
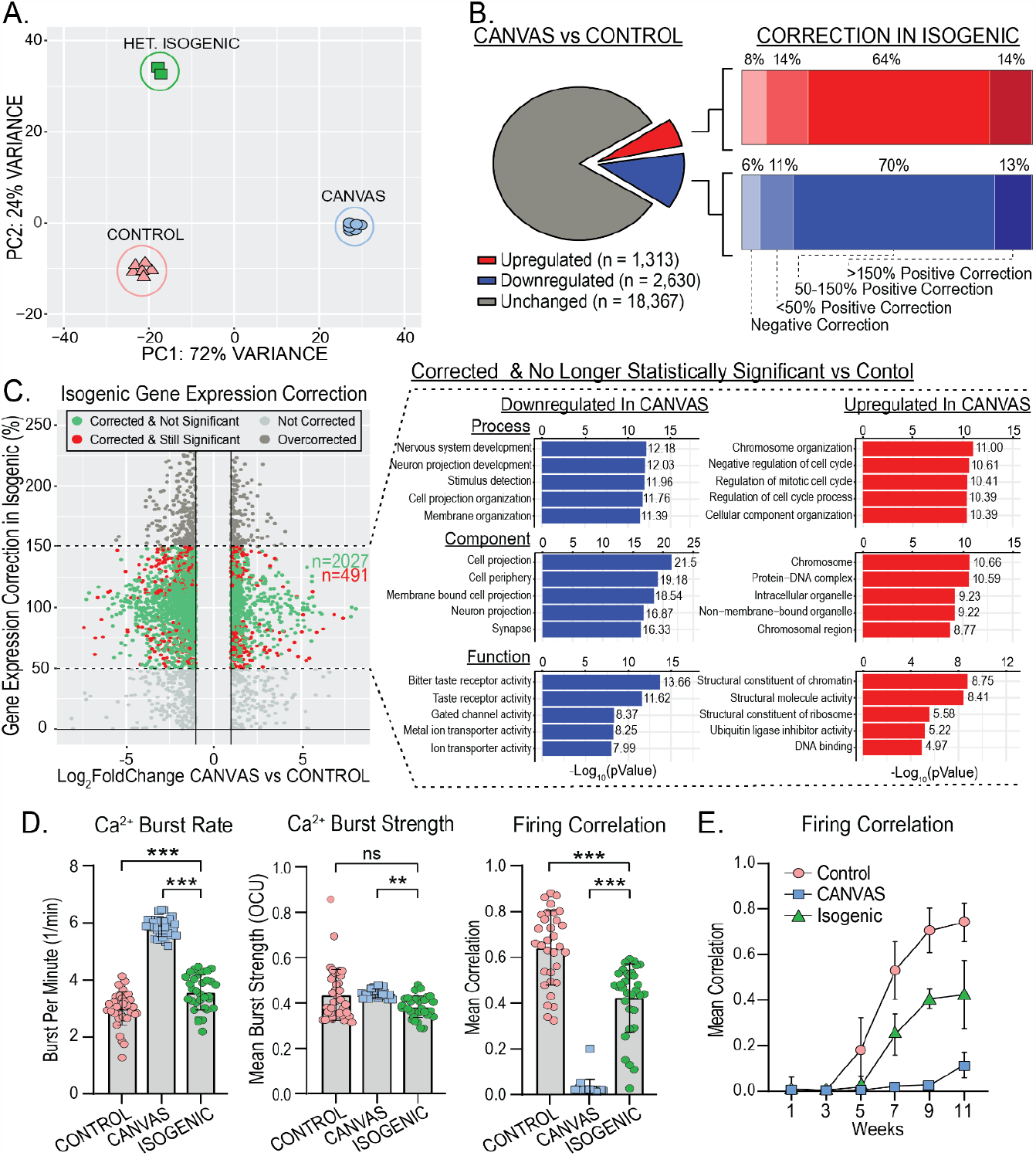
Heterozygous isogenic correction of CANVAS neurons corrects transcriptomic and synaptic functional deficits. **A**) Principal Component Analysis (PCA) of CANVAS (n=6), control (n=6), and CANVAS Heterozygous Isogenic (n=2) patient iPSC-derived neurons. **B**) Schematic illustrating the total number of genes dysregulated (up & down) in CANVAS vs control (left), with the percentage of these up- or down-dysregulated genes that show negative correction, partial correction, or full correction of expression upon heterozygous isogenic correction of CANVAS patient iPSC-derived neuron line. **C**) Scatter plot of Log_2_FoldChange (CANVAS vs control) vs Gene expression correction per gene in the heterozygous isogenic patient iPSC-derived neurons (left), and Gene Ontology (GO) pathway analysis of the top 5 up/downregulated Biological Process, Cellular Component, and Molecular Function for the genes that show 50-150% gene expression in Isogenic Correction vs CANVAS and are non-statistically significant in Isogenic vs control conditions. **D**) Analysis of Ca^2+^ imaging metrics for control (n=3), CANVAS (n=3), and Heterozygous Isogenic (n=1) patient iPSC-derived neurons. Burst Rate (F(2, 165) = 279.4, P<0.0001), Burst Strength (F(2, 165) = 4.034, P=0.019), Firing Correlation (F(2, 165) = 185.9, P<0.0001). Each data point represents the mean of ∼1000-3000 active cells per well (Supplementary Figure 8). Data were analyzed by one-way ANOVA with Sidak’s post-hoc multiple comparison tests. **E**) Mean firing correlation of control (n=3), CANVAS (n=3), and heterozygous isogenic (n=1) patient iPSC-derived neurons across 10 weeks of differentiation from week 1 to week 11. Error = SD.

### Synaptic gene expression is corrected upon heterozygous correction of *RFC1* repeat expansion

To determine the impact of the AAGGG/CCCTT repeat expansion in CANVAS patient iPSC-derived neurons, we generated an isogenic line with a monoallelic deletion of the repeat and compared its gene expression to both CANVAS and control iPSC-derived neuronal lines. Surprisingly, the heterozygous isogenic line almost completely corrected across PC1 that separates CANVAS from control neuronal samples, while variance along PC2 was increased substantially, indicating partial correction of some additional CANVAS-associated variance to control and the emergence of a new source of variance after heterozygous deletion of the AAGGG/CCCTT repeat expansion (Figure 6A, Supplementary Figure 10A-B). This large-scale correction of the CANVAS associated transcriptomic signature is also visible by heatmap analysis illustrating that the heterozygous isogenic patient iPSC-derived neurons exhibit a global gene expression pattern more similar to that of control than CANVAS (Supplementary Figure 10C).

In total, 64% of the CANVAS upregulated genes and 70% of the CANVAS down-regulated genes exhibited at least a 50% correction in expression upon monoallelic deletion of the AAGGG/CCCTT repeat expansion in the CANVAS patient 1 line (Figure 6B), and 80.5% of these genes (2027:491) were no longer significantly different compared to controls (Figure 6C, Green). Importantly, the GO-terms of these gene expression corrected genes correlate with many of the top dysregulated pathways in CANVAS, including multiple synapse-associated processes and functions (Figure 6C, right, Supplementary Figure 10D). Within these pathways, the key synaptic genes found to be most downregulated in CANVAS (Figure 4E) show significant or almost complete restoration in the isogenic neurons compared to control (Supplementary Figure 10E), with *CAMKIIB, GAP43, HOMER1, NEURL1*, and *SYP* showing the greatest restoration in the isogenic line.

### Dysregulated synaptic activity is rescued by heterozygous correction of *RFC1* repeat expansion

We next analyzed spontaneous synaptic activity upon heterozygous isogenic correction in comparison to both control and CANVAS patient iPSC-derived neurons (Figure 6D). Heterozygous isogenic neurons exhibited modest but significant improvements in basal calcium intensity and burst duration compared to CANVAS neurons (Supplementary Figure 9B), as well as large and significant improvements in neuronal burst rate synchronized firing (Figure 6D-E), with an average network correlation of 42% compared to 70% in controls. While these metrics don’t indicate complete recovery of deficits observed in CANVAS versus control, the significant trend towards recovery in all metrics in conjunction with the observed correction in gene expression upon deletion of a single expanded allele in CANVAS neurons suggests a repeat-dependent mechanism of CANVAS pathogenesis.

### *RFC1* reduction does not mimic CANVAS-associated transcriptomic or functional deficits

If loss of RFC1 function is central to CANVAS pathogenesis, then targeted loss of RFC1 should recapitulate defects observed in CANVAS patient neurons. To test this hypothesis, we conducted transcriptomic- and functional neuronal analyses of control neurons after sustained (∼10 weeks) knockdown of *RFC1* in comparison to CANVAS neurons. Treatment with RFC1 shRNA lentivirus led to undetectable levels of RFC1 protein (Figure 7A), and normalized *RFC1* gene counts across experimental conditions showed efficient knockdown of *RFC1* transcripts with ∼63% reduction in comparison to control neurons and ∼75% reduction in comparison to CANVAS neurons (Figure 7B, left). Compared to control neurons expressing a non-targeting shRNA, *RFC1* knockdown led to statistically upregulated expression of only 0.5% (235) of detected genes and statistically downregulated expression of only 0.23% (103) of detected genes in iPSC-derived neurons. PCA clustering showed no overlap between *RFC1* shRNA treated iPSC-derived neurons and CANVAS neurons and instead showed tight clustering with non-targeting shRNA expressing controls (Figure 7B, right, Supplementary Figure 11A-B), indicating that *RFC1* knockdown in control neurons does not induce CANVAS-like transcriptomic alterations. Further, volcano plot of *RFC1* knockdown vs control neurons (Figure 7C) indicates less dysregulation in gene expression when compared to CANVAS vs control (Figure 4A), with a preference for increased expression for a small number of transcripts and a reduction in *RFC1* (circled). Similarly, none of the key synaptic genes found to be highly downregulated in CANVAS (Figure 4E) showed significant dysregulation upon *RFC1* knockdown (Supplementary Figure 11C).

**Figure 7.**
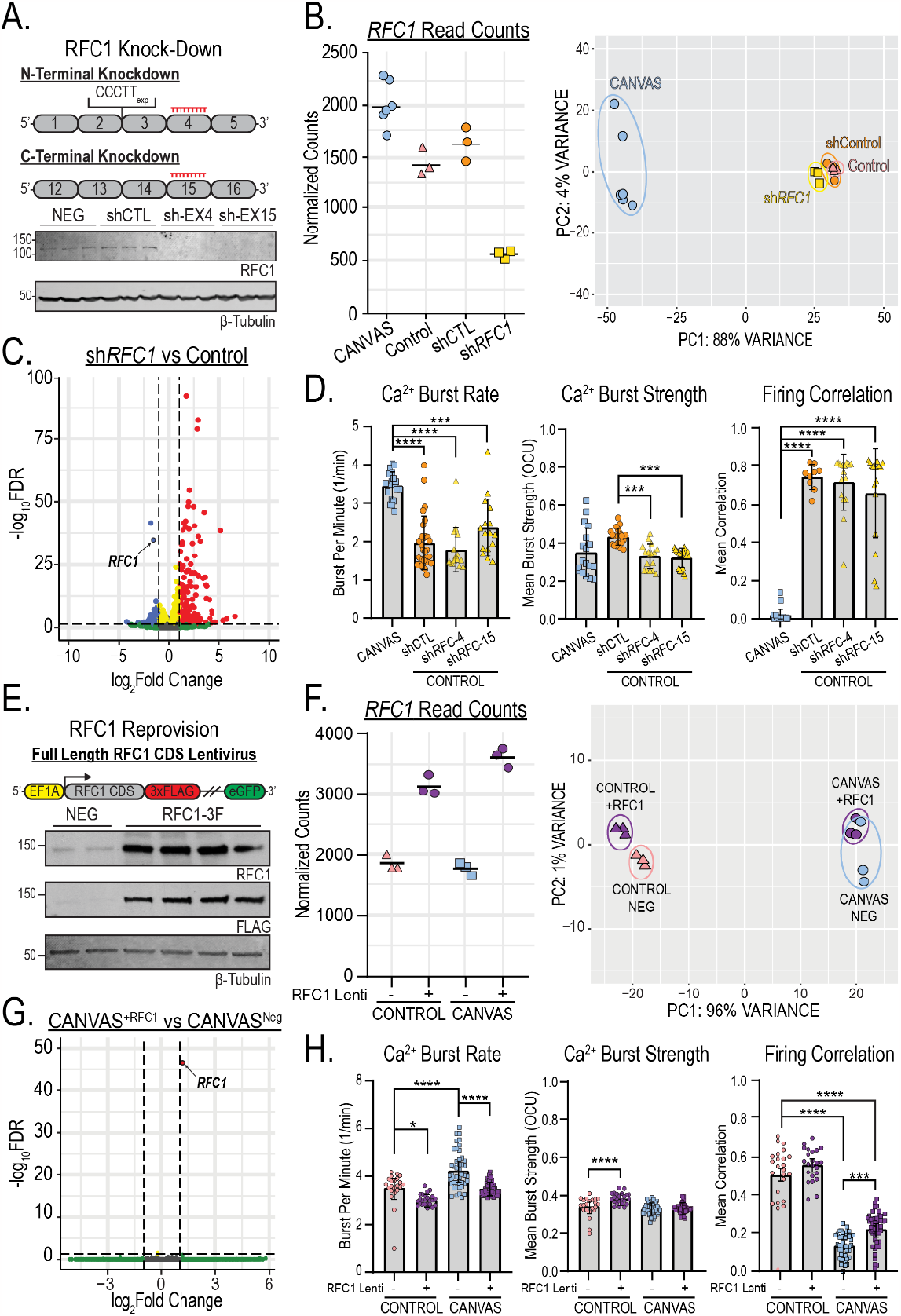
RFC1 knockdown or reprovision fail to recapitulate or correct transcriptomic and synaptic deficits in CANVAS patient-iPSC derived neurons. **A**) Schematic outlining approach to knockdown of either the N- or C-terminus of RFC1 by shRNA lentiviral transduction (Top), and analysis of RFC1 expression from control iPSC-derived neurons (GTX129291, 1:1000) indicating successful knockdown of RFC1 (Bottom). **B**) Normalized read counts for RFC1 transcripts in CANVAS, control iPSC-derived neurons as well as control iPSC-derived neurons treated with either control or anti-RFC1 shRNA lentivirus (left), and Principal Component Analysis (PCA) of CANVAS (n=6), control (n=3), control mock-treated (n=3), and control sh*RFC1* treated (n=3) patient iPSC-derived neurons. **C**) Volcano plot of -Log_10_FDR vs Log_2_(Fold Change) for RFC1 knockdown vs control, RFC1 labelled. **D**) Analysis of Ca^2+^ imaging metrics for CANVAS (n=3) and control (n=3) patient iPSC-derived neurons treated with sh*Control* or sh*RFC1* exon4/exon15 lentiviruses. Burst Rate (F(3, 78) = 29.6, P<0.0001), Burst Strength (F(3, 78) = 8.265, P<0.0001), Firing Correlation (F(3, 78) = 100.6, P<0.0001). **E**) Schematic outlining the approach of RFC1 overexpression in CANVAS patient iPSC-derived neurons by lentiviral transduction (Top), and analysis of RFC1 expression in patient iPSC-derived neurons upon lentiviral transduction (Bottom). **F**) Normalized read counts for RFC1 transcripts in CANVAS and control iPSC-derived neurons transduced with either full-length RFC1 CDS lentivirus or control lentivirus (n=3/group) (left), and Principal Component Analysis (PCA) of CANVAS and control iPSC-derived neurons transduced either full-length RFC1 CDS lentivirus or control lentivirus (n=3/group) (right). **G**) Volcano plot of -Log_10_FDR vs Log_2_(Fold Change) for CANVAS patient-derived neurons transduced with either full-length RFC1 CDS lentivirus or control lentivirus (n=3/group), RFC1 labelled. **H**) Analysis of Ca^2+^ imaging metrics for control (n=3) and CANVAS (n=3) patient iPSC-derived neurons treated with control or RFC1-overexpression lentivirus. Burst Rate (F(3, 135) = 31.01, P<0.0001), Burst Strength (F(3, 135) = 16.74, P<0.0001), Firing Correlation (F(3, 135) = 147.3, P<0.0001). Firing Correlation two-way ANOVA treatment vs genotype: F(1,135) = 41.25, P<0.0001, F(1,135) = 36.64, P<0.0001 respectively. Each data point represents the mean of ∼1000-3000 active cells per well (Supplementary Figure 5). Data were analyzed by one-way ANOVA with Sidak’s post-hoc multiple comparison tests. Error = SD.

Gene set enrichment analysis (GSEA) of ranked genes from *RFC1* knockdown showed an enrichment for annotated RFC1-associated functions (Supplementary Figure 12). Gene ontology (GO) analysis for the most overrepresented cellular pathways within the transcripts dysregulated upon RFC1 knockdown, meanwhile, found that immune and developmental cellular processes were most significantly dysregulated (Figure 7C). There was no enrichment in synaptic-associated processes, in contrast to the significantly dysregulated pathways identified in CANVAS neurons vs control (Figure 4B, Supplementary Figures 8 & 11D).

To assess the impact of *RFC1* knockdown on neuronal function, we again used calcium imaging. Compared to control neurons, *RFC1* knockdown elicited no differences in basal calcium intensity, burst duration (Supplementary Figure 9C), neuronal burst rate or network firing correlation (Figure 7D) and the *RFC1* knockdown neurons remained highly significantly different from CANVAS neurons in these four metrics. The only metric that showed any similarities with the findings in CANVAS neurons was burst strength, where knockdown of *RFC1* induced a 24% reduction (Figure 7D, center, P<0.0001), comparable to the levels seen in CANVAS neurons (CANVAS – shRNA1: P=0.915, CANVAS – shRNA2: P=0.786).

### Reprovision of RFC1 does not correct CANVAS-associated transcriptomic or functional deficits

To determine if reprovision of RFC1 can correct phenotypic deficits observed in CANVAS patient neurons, we conducted transcriptomic- and functional neuronal analyses of control and CANVAS iPSC-derived neurons after sustained (∼10 weeks) expression of either GFP lentivirus or a full-length RFC1 lentivirus construct with a C-terminal 3x FLAG tag and non-fused GFP (Figure 7E). Transduction of this lentivirus significantly boosted RFC1 expression in patient iPSC-derived neurons (Figure 7E), with a ∼50-75% increase in detected *RFC1* transcripts compared to neurons treated with the GFP control lentivirus only (Figure 7F, left). PCA clustering showed no overlap of RFC1 overexpression CANVAS neurons with control neurons and instead showed tight clustering between genotypes independent of treatment condition (Figure 7F, right), indicating that *RFC1* overexpression fails to correct CANVAS neuronal transcriptomic alterations. Further, volcano plot of differential gene expression in CANVAS neurons overexpressing RFC1 compared to those expressing a control GFP lentivirus show that sustained reprovision of RFC1 to CANVAS neurons induces effectively no transcriptomic changes, with only *RFC1* found to be differentially expressed between these conditions (Figure 7G). Similarly, comparison of CANVAS neurons overexpressing RFC1 with control neurons (Supplementary Figure 13A) exhibited identical differential gene expression patterns as was observed in Figure 4A. Overexpression of RFC1 in control neurons also triggered minimal changes in global gene expression (Supplementary Figure 13B), and heatmap profiles of global gene expression across all conditions indicate that overexpression of RFC1 does not correct the transcriptomic deficits observed in CANVAS, and elicits negligible effects within control neurons (Supplementary Figure 13C).

When analyzing the functional phenotypes observed for CANVAS neurons, overexpression of RFC1 elicited no differences for metrics that exhibited only minor or no dysregulation in CANVAS compared to controls (Supplementary Figure 7D). Reprovision of RFC1 did reduce the abnormal firing burst rate in CANVAS neurons by 17.6%, which effectively corrected the difference between CANVAS and control neurons. However, RFC1 overexpression also induced a 13.9% reduction in burst rate within control neurons, suggesting that this effect is not specific to CANVAS neurons (Figure 7H, left). 2-way ANOVA analysis indicates that the effect sizes are comparable and that the CANVAS vs control genotype is not a driver of the effect (Treatment: F(1,135)=41.25, P<0.0001, Genotype: F(1,135)=36.64, P<0.0001). Similarly, RFC1 overexpression increased the firing correlation observed for CANVAS neurons (0.13 to 0.22, P=0.001). However, this correction was modest, and these CANVAS neurons remained significantly less correlated than control neurons (P<0.0001, Figure 7H, right).

## Discussion

With an estimated carrier frequency upwards of ∼6%^8,14^, non-reference AAGGG/CCCTT repeat expansions in *RFC1* are potentially significant contributors to neurologic disease, including both ataxia and sensory neuronopathy. Here we used patient-derived neurons, patient cells and tissues to directly assess how these repeats elicit toxicity. Our findings suggest a specific role for the repeat element in neuronal development and synaptic function that is largely independent of RFC1 expression and its canonical functions. These findings have important implications for therapy development in this currently untreatable disorder.

CANVAS arises from a polymorphic set of biallelic *RFC1* repeat expansion motifs comprising AAGGG, AAAGG, ACAGG, AAGGC, AGGGC, and AGAGG motifs in isolation or as heterozygous combinations^8,9,11,12^. Further, rare CANVAS patients harbor compound heterozygous monoallelic *RFC1* expansions combined with loss-of-function mutations that result in RFC1 haploinsufficiency from the non-expanded allele^17–19^. This, in combination with data indicating a recessive mode of inheritance, suggested to us and others that RFC1 loss-of-function is a pathogenic driver of CANVAS pathogenesis. However, we find that RFC1 expression is normal at both the mRNA and protein level in the context of biallelic expansions in differentiated human iPSC-derived neurons, with no changes in *RFC1* mRNA splicing, intron degradation, transcript isoform, or exon usage in CANVAS neurons compared to controls (Figure 3A-B, Supplementary Figure 4). Consistent with this, we observe no deficits in canonical RFC1 functions in either mitotic- and post-mitotic CANVAS patient-derived cell types. RFC1 plays a key role in both DNA replication and DNA damage repair^52^, but CANVAS patient-derived neurons do not accumulate DNA damage compared to controls, and they exhibit normal rates of recovery after UV-induced DNA damage (Figure 3, Supplementary Figure 6). Deficits in the rate compared to controls. CANVAS patient-derived neurons exhibit significant dysregulation in genes associated with pathways regulating synaptic of cell division and proliferation were observed in CANVAS patient fibroblasts (Figure 3C). However, these deficits were independent of RFC1 function, as they were neither replicated by RFC1 knockdown in control fibroblasts nor corrected by reprovision of RFC1 to CANVAS patient fibroblasts. Despite the clinical genetic data strongly suggesting an RFC1 loss-of-function in CANVAS and our own data showing gene alterations and signaling dysfunction in CANVAS patient iPSC-derived neurons, we find no evidence for a loss of canonical RFC1 function in multiple cell types, pointing towards a more nuanced etiology driving CANVAS pathogenesis at the *RFC1* locus.

Repeat associated gain-of-function mechanisms, such as the formation of RNA foci and the presence of ubiquitinated inclusions of RAN translated repeat polypeptides, occur in many repeat expansion neurologic diseases^31,37–39,55^. Yet, postmortem and cell-based analyses performed to date have not demonstrated any of these pathological features in CANVAS^7,8^. We detected sense strand AAGGG repeat-derived pentapeptide repeat peptides in cerebellar granule cells in CANVAS patient, but not control, brains (Figure 2, Supplementary Figure 3). We did not detect these proteins or repeat RNA foci in patient iPSC-derived neurons and overexpression of these AAGGG repeats in isolation did not induce neuronal toxicity (Figure 2, Supplementary Figure 2). It is therefore unclear what role – if any-these translated pentapeptide repeats play in the neuropathogenic cascades that precede CANVAS symptom onset. CANVAS patient iPSC-derived neurons showed significant deficits in synaptic gene expression at both the mRNA and protein level structure and organization, regulation of chemical synaptic transmission, and expression of ion channels localized to both the pre- and post-synaptic membranes (Figures 4-5, Supplementary Figure 8). This deficit correlates with phenotypic deficits in synaptic activity (Figures 5-7). While synaptic dysfunction and loss of synaptic connections occurs in numerous neurodegenerative diseases, such as Huntington Disease (HD)^56,57^, *C9orf72* FTD/ALS^58,59^, and Alzheimer Disease (AD)^60,61^, the molecular steps preceding this deficit in synaptic gene expression and associated synaptic dysfunction in CANVAS patient neurons is unclear. Intriguingly, monoallelic deletion of the AAGGG expansion significantly corrected both synaptic gene expression defects and synaptic signaling dysfunction in CANVAS neurons (Figure 6). As this isogenic deletion of a single AAGGG expansion does not alter RFC1 expression or neuronal response and recovery after UV induced DNA damage (Supplementary Figure 6), this rescue in synaptic gene expression and neuronal connectivity instead appears to be dependent on the repeat element itself.

If dysfunction in RFC1 protein expression or activity was the cause of CANVAS phenotypes, then we would predict that synaptic dysfunction and gene expression alterations in CANVAS patient-derived neurons would be replicated by knockdown of *RFC1* in control neurons. However, sustained (∼10 weeks) knockdown of *RFC1* with two independent shRNAs in control iPSC-derived neurons elicits no deficits in synaptic function or gene expression changes akin to those seen in CANVAS neurons (Figure 7, Supplementary Figures 9, 11, 12). Similarly, if RFC1 loss impacts CANVAS disease relevant phenotypes, then prolonged reprovision of RFC1 protein into CANVAS patient iPSC-derived neurons should rescue the functional synaptic phenotypes observed. Yet, we observed no correction of CANVAS neuronal transcriptomic signatures after RFC1 reprovision. Moreover, while minor changes were observed in CANVAS neuronal activity with exogenous RFC1 expression, these improvements were both modest and non-specific, with similar effects in control and CANVAS neurons. CANVAS neurons remained highly dysregulated even when expressing RFC1 (Figure 7, Supplementary Figures 9, 11).

In summary, our studies provide strong support for a repeat-dependent mechanism of neuronal dysfunction in CANVAS that operates outside of the canonical functions of RFC1. These findings stand in contrast to clinical genetic studies pointing towards an RFC1 loss-of-function mechanism as a central contributor to the molecular etiology of CANVAS and suggest that replacing or boosting canonical RFC1 function would be ineffective as a therapeutic approach in this condition. Moreover, the mechanism by which these repeats act does not fit into known repeat-associated gain-of-function and loss-of-function boxes previously defined for other repeat expansion disorders. Instead, our findings suggest that these non-reference repeats act through an as-yet undefined novel molecular mechanism that will likely have relevance beyond this condition.

## Supplementary Figures

**Supplementary Figure 1.**
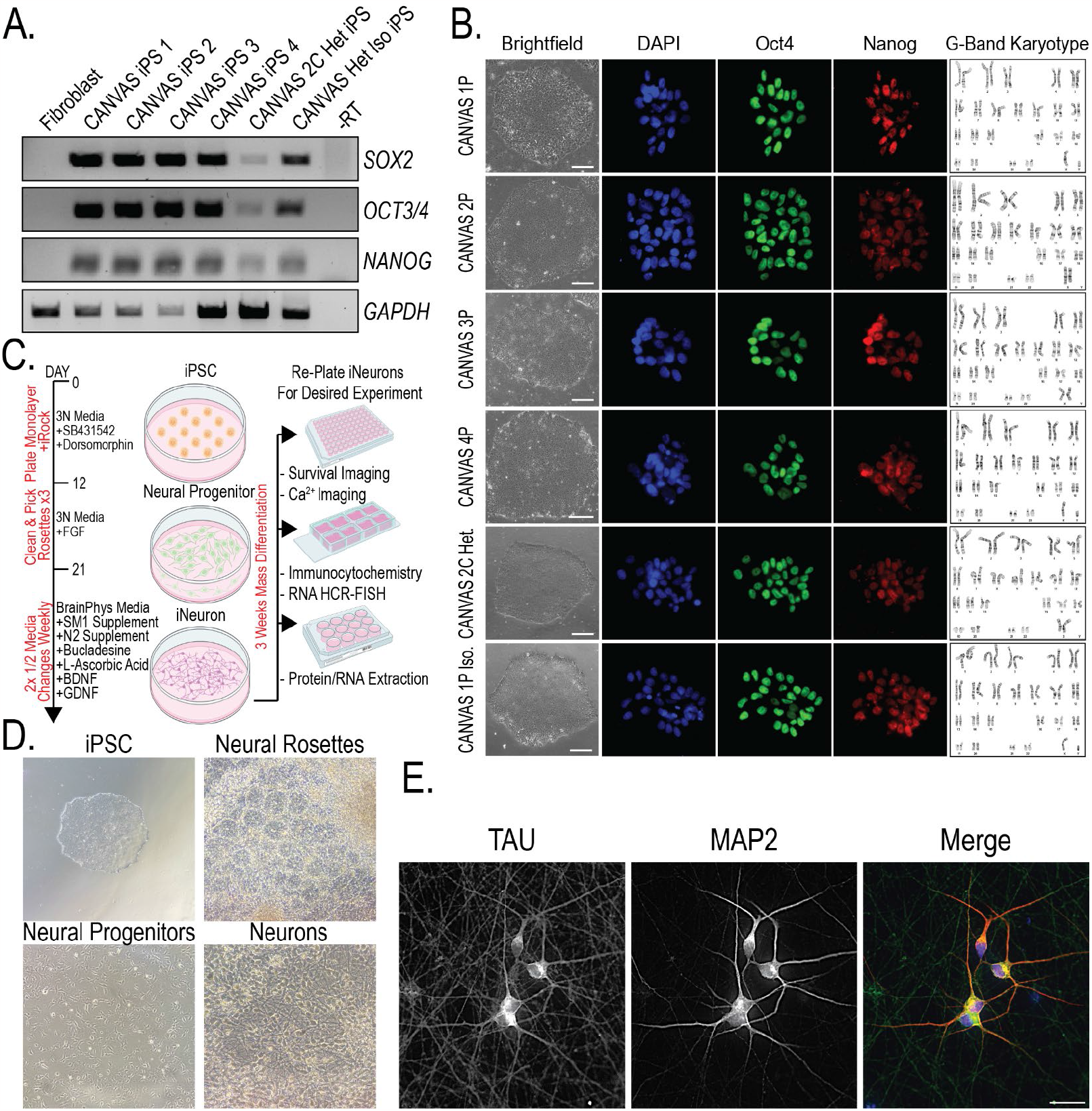
Generation, characterization, and differentiation of CANVAS patient iPSC-derived neurons. **A**) RT-PCR of CANVAS patient derived iPSCs for the pluripotency mRNA markers *SOX2, OCT3*/*4*, and *NANOG*. **B**) G-Band Karyotype analysis (WiCell) and Immunocytochemistry of CANVAS patient iPSCs for the pluripotency markers OCT4 and NANOG, scale = 25 μm. **C**) Schematic outlining the process of differentiating patient-derived iPSCs to neural progenitor cells and glutamatergic neurons by dual-SMAD inhibition for experimentation. **D**) Brightfield images of the stages of neuronal differentiation from patient-derived iPSCs, showing iPSCs, neural rosettes, neural progenitor cells, mass-differentiated neuronal cells and re-plated neuronal cells for experimentation. **E**) Immunocytochemistry of a representative patient iPSC-derived neurons stained for the neuronal markers Tau and MAP2. Scale = 25 μm.

**Supplementary Figure 2.**
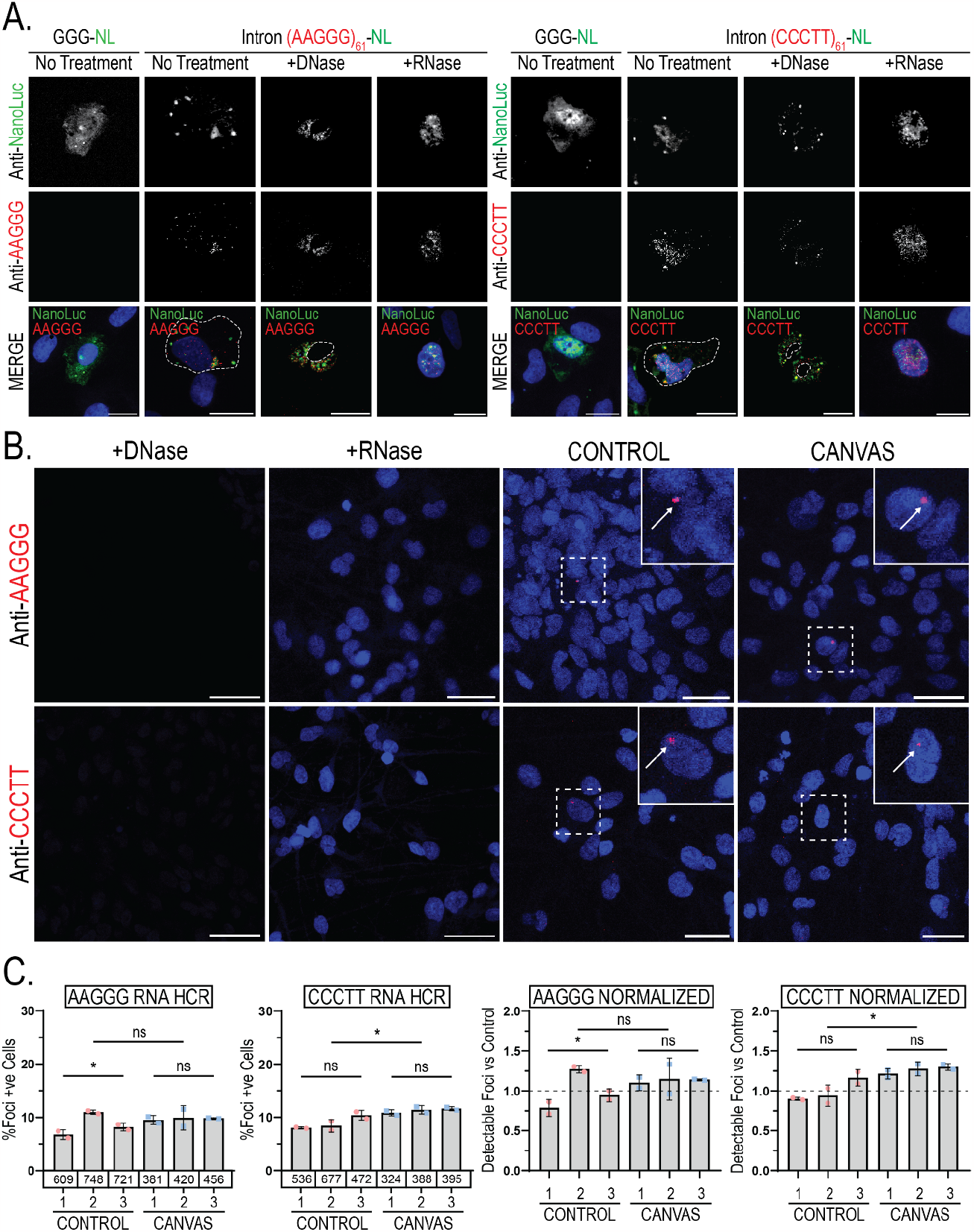
RNA HCR probe validation and analysis of RNA foci formation in CANVAS patient and control iPSC-derived neurons. **A**) Confocal images of HEK293 cells transfected with control GGG-NL plasmid or plasmid expressing intronic AAGGG or CCCTT expansion followed by RNA HCR-FISH for AAGGG and CCCTT RNAs. Cells were fixed and treated with either anti-Nanoluc, anti-AAGGG, or anti-CCCTT fluorescent probes after no treatment, DNase, or RNase treatment to assess probe specificity. Scale = 10 μm. **B**) Confocal Images of representative CANVAS and control patient iPSC-derived neurons after RNA HCR-FISH utilizing either anti-AAGGG or anti-CCCTT fluorescent probes to assess sense or antisense RNA foci formation. Scale = 10 μm. **C**) Quantification of foci positive neurons for control (n=3) and CANVAS (n=3) patient iPSC-derived neurons with total n-numbers of neuronal cells analyzed per cell line indicated. AAGGG (F(5, 12) = 3.619, P=0.074), CCCTT (F(5, 12) = 8.293, P=0.011). Data were analyzed by one-way ANOVA with Sidak’s post-hoc multiple comparison tests. Error = SD.

**Supplementary Figure 3.**
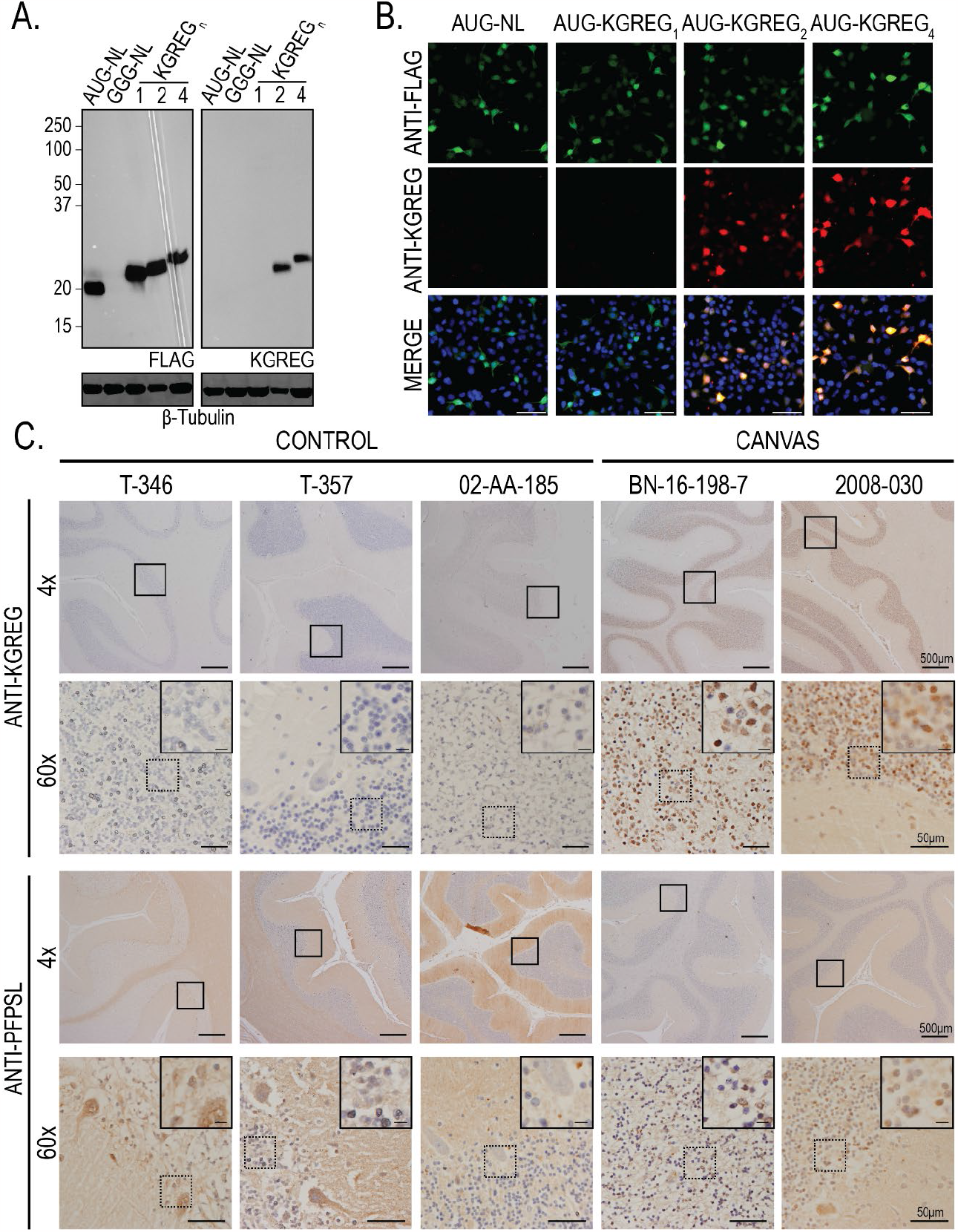
Analysis of repeat-derived peptides using KGREG & PFPSL antibodies. **A**) Analysis of lysates from HEK293 cells transfected with control plasmid or plasmids expressing 1x, 2x, and 4x KGREG FLAG-tagged plasmids using anti-FLAG M2 (1:1000) and anti-KGREG (1:100) antibodies. **B**) ICC of HEK293 cells transfected with control plasmid or plasmids expressing 1x, 2x, and 4x KGREG FLAG-tagged plasmids using anti-FLAG M2 (1:100) and anti-KGREG (1:100) antibodies, scale = 25 μm. **C**) Immunohistochemistry of control (n=3) and RFC1 expansion CANVAS (n=2) patient post-mortem cerebellar vermis tissue stained with sense anti-KGREG or antisense anti-PFPSL antibodies (1:100, acid AR). Scale = 500 μm (4x), 50 μm (60x) and 20 μm (inset).

**Supplementary Figure 4.**
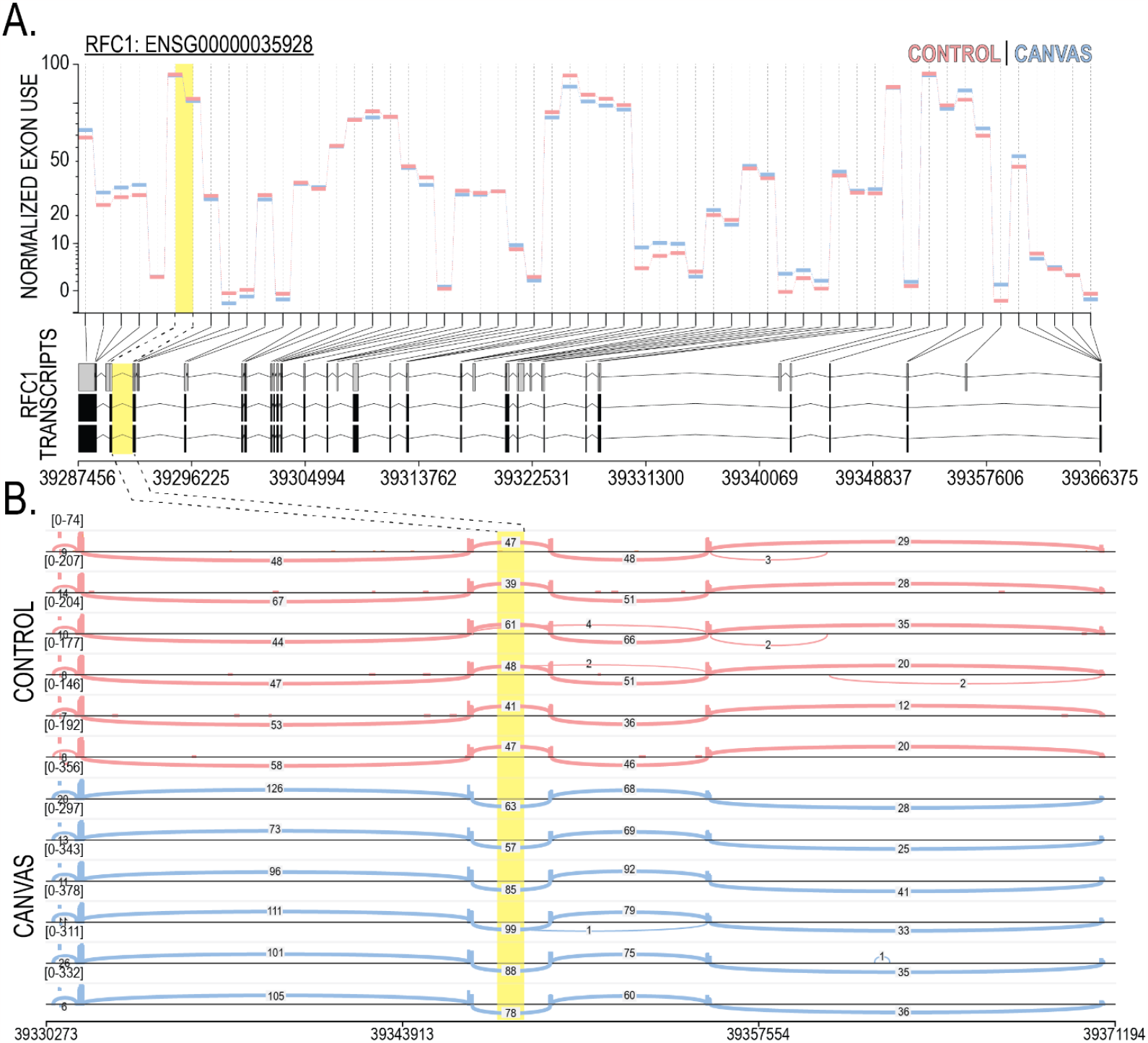
Analysis of *RFC1* isoform and alternative exon usage RNA short-read sequencing. **A**) Normalized exon usage for *RFC1* mRNA transcripts in CANVAS patient iPSC-derived neurons, analyzed by DEXSeq of paired-end RNASeq reads. **B**) Sashimi plot of *RFC1* exons 1-5 in CANVAS patient iPSC-derived neurons indicating no skipping of exons flanking the intron 2 repeat expansion region (highlighted yellow).

**Supplementary Figure 5.**
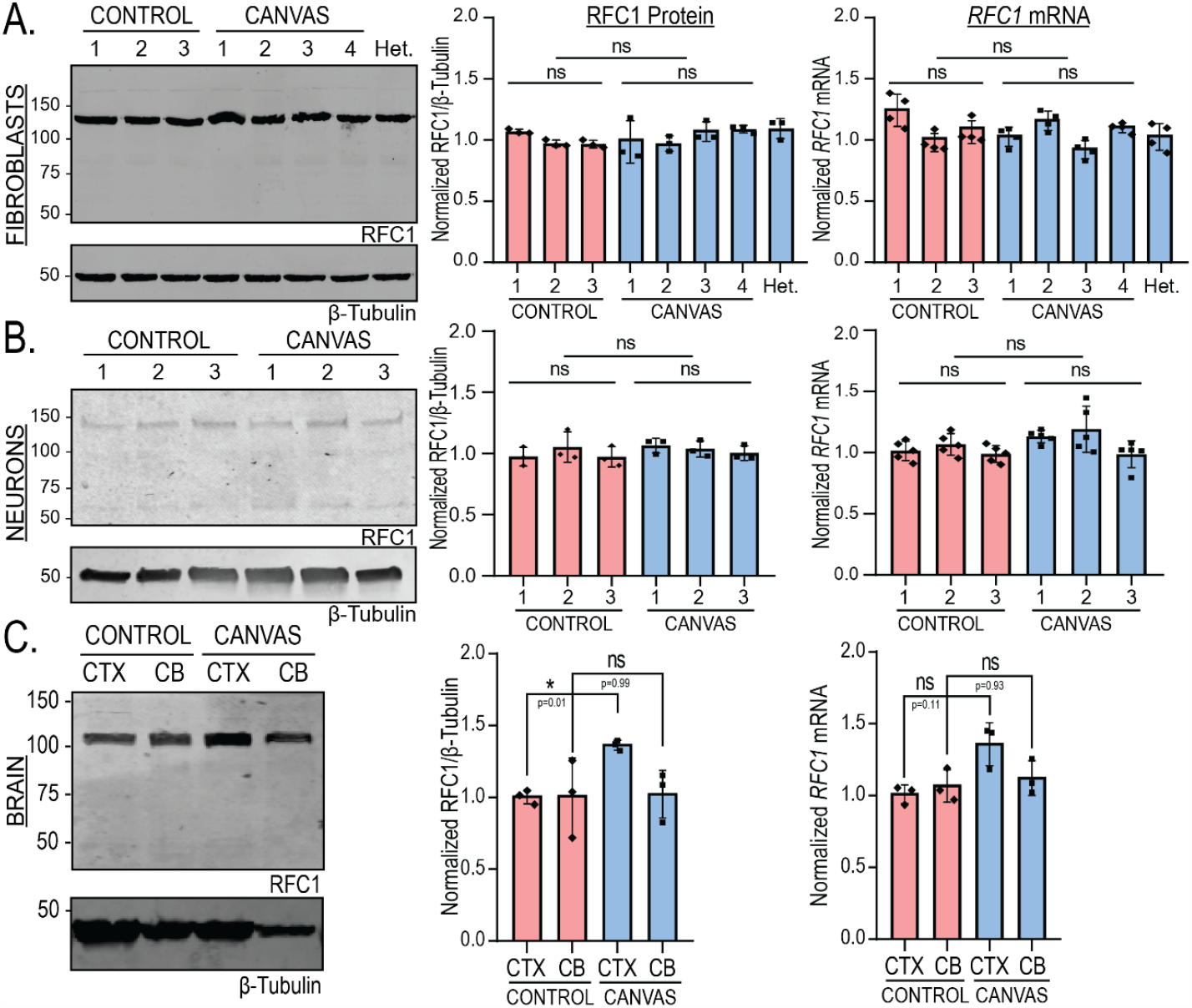
RFC1 protein and mRNA expression is maintained in AAGGG expansion contexts. **A**) Analysis of RFC1 expression (**left**), quantification of normalized RFC1 expression (**center**, F(7,24) = 1.592, P=0.208), and quantification of normalized RFC1 mRNA expression (**right**, F(7,24) = 4.944, P=0.0014) from CANVAS (n=4) and control (n=3) patient-derived fibroblasts. **B**) Analysis of RFC1 expression (**left**), quantification of normalized RFC1 expression (**center**, F(5,18) = 0.707, P=0.629)), and quantification of normalized RFC1 mRNA expression (**right**, F(5,24) = 3.029, P=0.029) from CANVAS and control (n=3) patient iPSC-derived neurons. **C**) Analysis of RFC1 protein (**left**), quantification of normalized RFC1 protein expression (**center**, F(3,8) = 2.010, P=0.191) from CANVAS (n=1) and control (n=1) patient post-mortem brain tissue, and quantification of normalized RFC1 mRNA expression (**right**, F(3,8) = 5.171, P=0.0281). N = 3 biological replicates. Data were analyzed by one-way ANOVA with Sidak’s post-hoc multiple comparison tests. Error = SD.

**Supplementary Figure 6.**
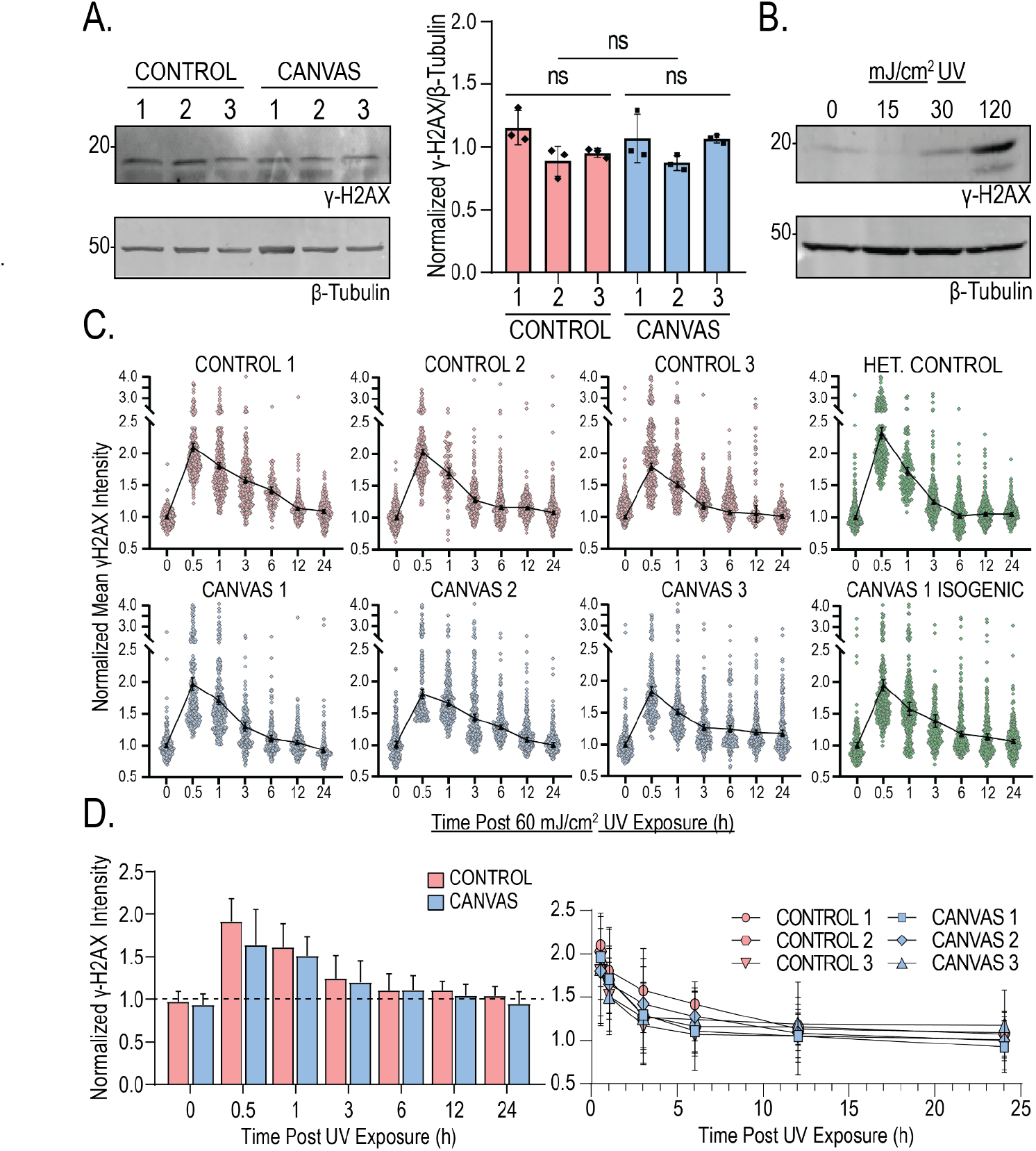
Analysis of DNA damage accumulation and recovery in patient iPSC-derived neurons. **A**) Analysis of γ-H2AX expression (**left**), and quantification of normalized γ-H2AX levels (**right**, F(5, 18) = 2.962, P=0.057) from CANVAS (n=3) and control (n=3) patient iPSC-derived neurons. Data were analyzed by one-way ANOVA with Sidak’s post-hoc multiple comparison tests. N = 3 biological replicates. **B**) Analysis of γ-H2AX levels in control iPSC-derived neurons exposed to 0, 15, 30, and 120 mJ/cm^2^ UV irradiation. **C**) Quantification of mean γ-H2AX staining in control (n=3), CANVAS (n=3), heterozygous *RFC1* expansion (n=1), and CANVAS heterozygous isogenic corrected (n=1) patient iPSC-derived NeuN+ neuronal nuclei over a 24h period after 60 mJ/cm^2^ UV exposure. **D**) Comparison of control (n=3) and CANVAS (n=3) γ-H2AX levels after 60 mJ/cm^2^ UV exposure (left) and comparison of first derivative rates of γ-H2AX decline (right, F(5, 30) = 0.033, P=0.999). Error = SD.

**Supplementary Figure 7.**
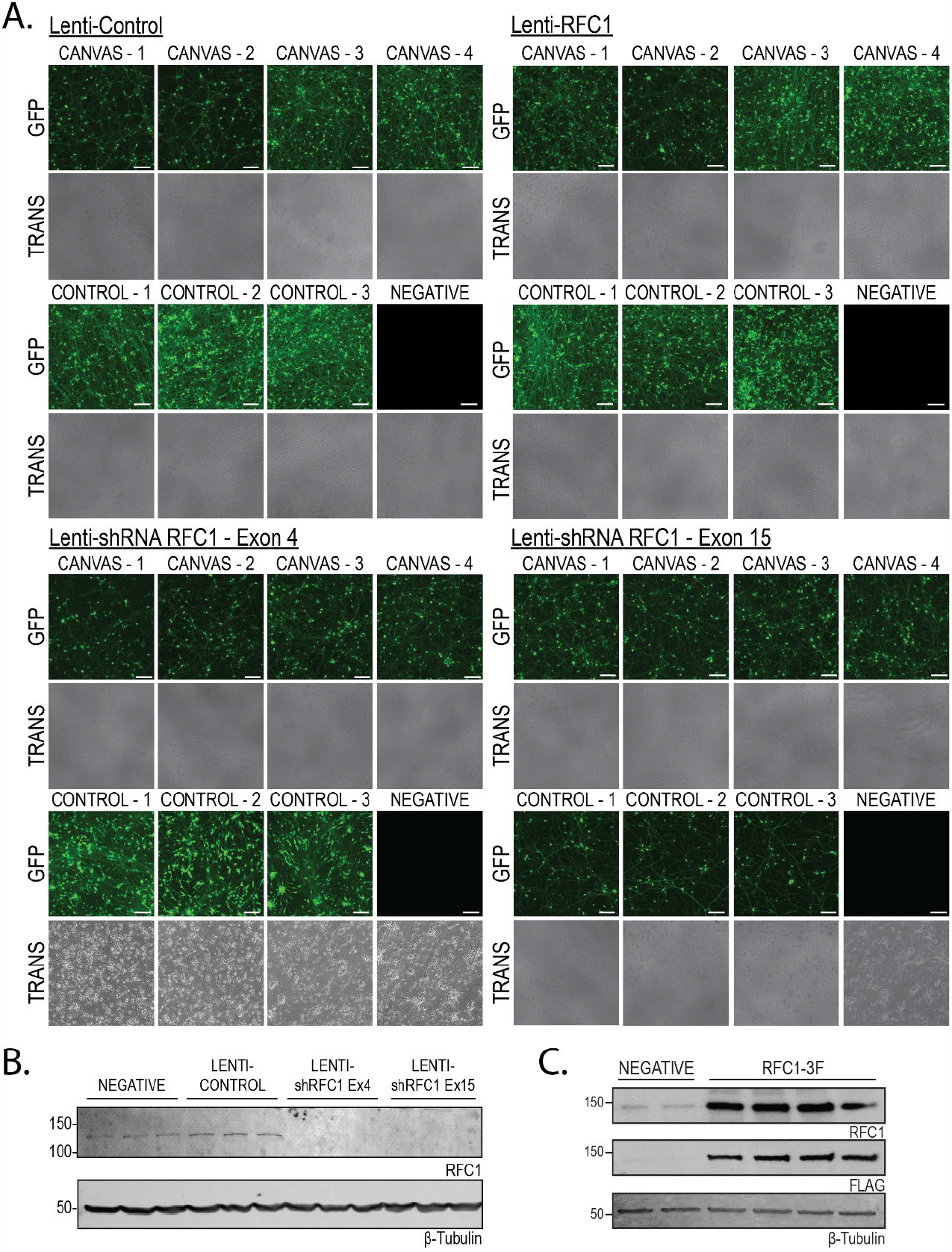
Efficiency of lentiviral transduction in CANVAS patient and control iPSC-derived neurons. **A**) Brightfield and GFP fluorescence images of CANVAS (n=4) and control (n=3) patient iPSC-derived neurons transduced with control lentivirus, RFC1 overexpression lentivirus, or lentiviruses encoding shRNAs to knockdown RFC1 at exon4 or exon15. Scale = 50 μm. **B-C**) Analysis of RFC1 expression (GTX129291 1:1000) upon shRNA RFC1 lentivirus treatment (**B**) or RFC1 overexpression lentivirus treatment (**C**) to assess for lentivirus efficiency.

**Supplementary Figure 8.**
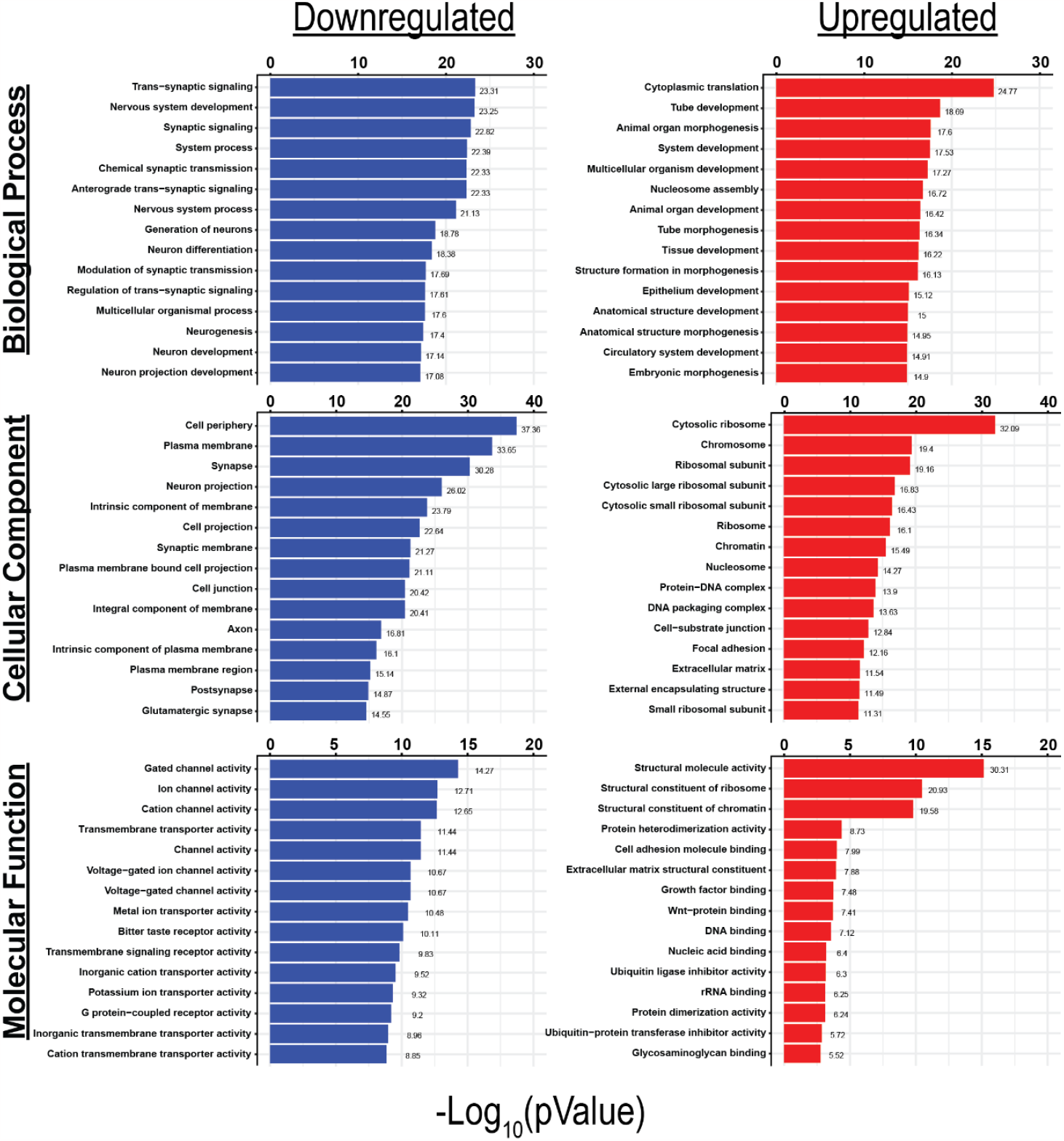
Gene ontology (GO) analysis of dysregulated transcripts detected in CANVAS patient vs control iPSC-derived neurons. Full Gene Ontology (GO) pathway analysis of the top up and downregulated Biological Process, Cellular Component, and Molecular Functions in CANVAS patient vs control iPSC-derived neurons. N = 6 biological replicates from 3 individual CANVAS and control patients.

**Supplementary Figure 9.**
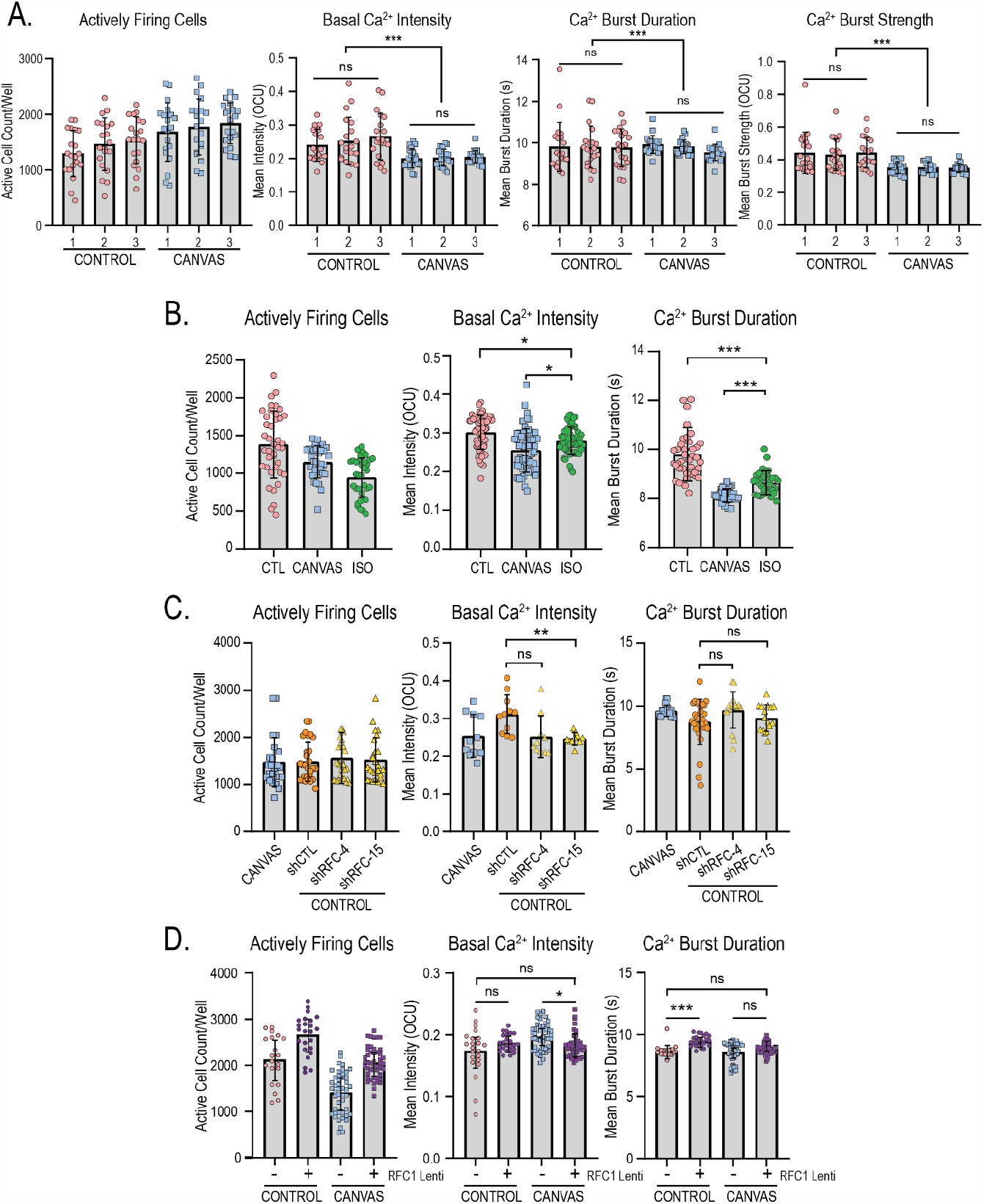
Calcium imaging metrics from CANVAS patient and control iPSC-derived neurons. **A)** Analysis of Ca^2+^ imaging metrics for control (n=3) and CANVAS (n=3) patient iPSC-derived neurons. Basal Intensity (F(5, 114) = 7.075, P<0.0001), Burst Duration (F(5, 114) = 0.5371, P=0.745), Burst Strength (F(5, 114) = 7.573, P<0.0001). **B**) Analysis of Ca^2+^ imaging metrics for control (n=3), CANVAS (n=3), and Heterozygous Isogenic (n=1) patient iPSC-derived neurons. Basal Intensity (F(2, 165) = 14.31, P<0.0001), Burst Duration (F(2, 101) = 48.79, P<0.0001). **C**) Analysis of Ca^2+^ imaging metrics for CANVAS (n=3) and control (n=3) patient iPSC-derived neurons treated with shControl or shRFC1 exon4/exon15 lentiviruses. Basal Intensity (F(3, 78) = 4.279, P=0.011), Burst Duration (F(3, 78) = 2.520, P=0.0641). **D**) Analysis of Ca^2+^ imaging metrics for control (n=3) and CANVAS (n=3) patient iPSC-derived neurons treated with control or RFC1-overexpression lentivirus. Basal Intensity (F(3, 135) = 5.621, P=0.0012), Burst Duration (F(3, 135) = 14.42, P<0.0001). Each data point represents the mean of ∼1000-3000 active cells per well. Data were analyzed by one-way ANOVA with Sidak’s post-hoc multiple comparison tests. N= 3 biological replicates, error = SD.

**Supplementary Figure 10.**
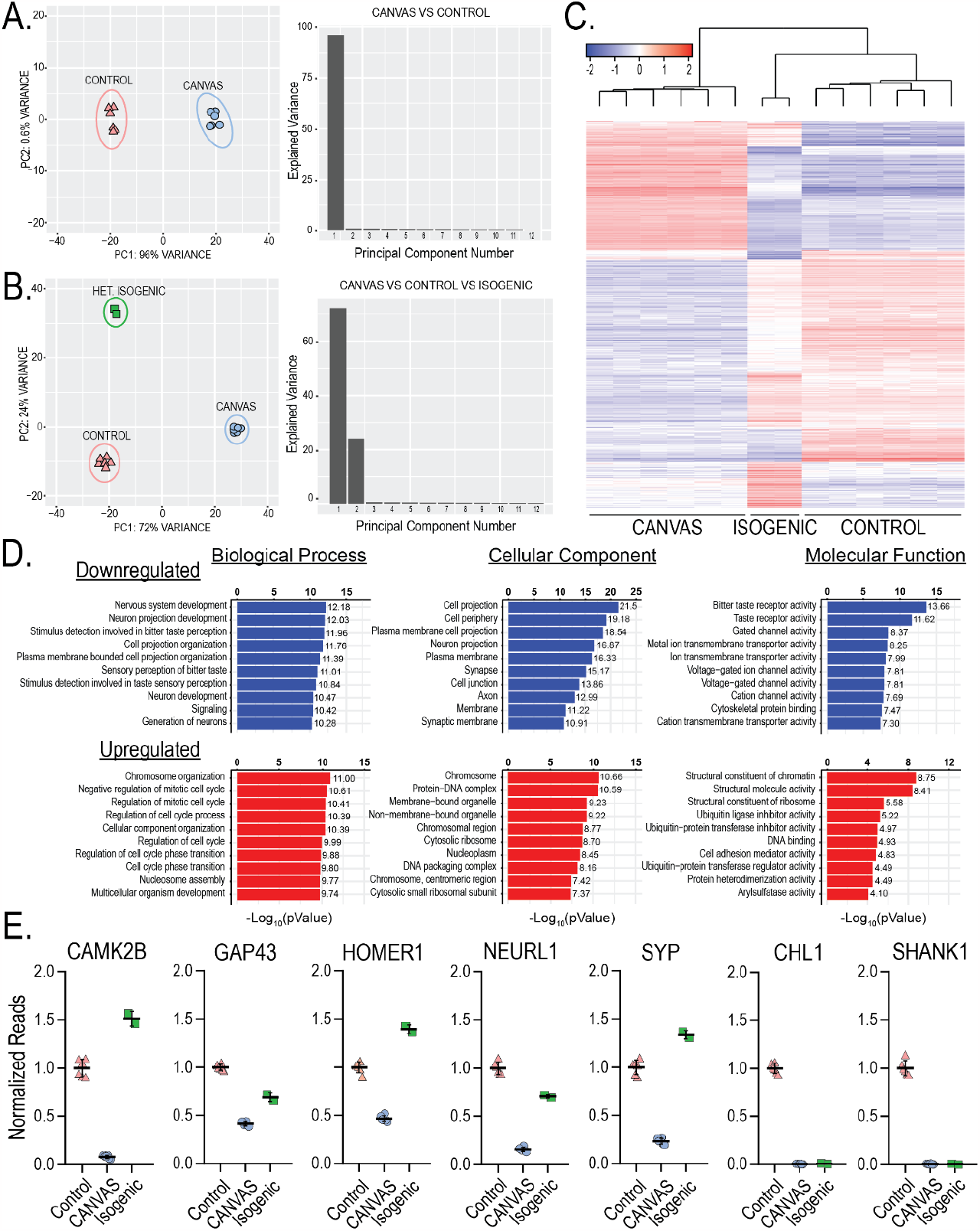
Supplemental data of transcriptomic analyses from control, CANVAS, and heterozygous isogenic correction patient iPSC-derived neurons. **A)** Principal component analyses of CANVAS (n=6) vs control (n=6) patient iPSC-derived neurons (left) with scree plot of explained variance with principal component axis of variance per sample shown for PC1 and PC2 (right). **B**) CANVAS (n=6), control (n=6), and Heterozygous Isogenic (n=2) patient iPSC-derived neurons (left) with principal component axis of variance per sample shown for PC1 and PC2 (right). **C**) Heatmap of normalized expression for the top 1000 genes differentially expressed in CANVAS, control, and CANVAS isogenic patient iPSC-derived neurons. **D**) Gene Ontology (GO) pathway analysis of the top up/downregulated Biological Process, Cellular Component, and Molecular Function for the genes that showed significant expression correction in heterozygous isogenic iPSC-derived neurons compared to CANVAS/control. **E**) Normalized read counts for *CAMK2B, GAP43, HOMER1, NEURL1, SYP, CHL1*, and *SHANK1* from control (n=6), CANVAS (n=6), and isogenic (n=2) iPSC-derived neurons.

**Supplementary Figure 11.**
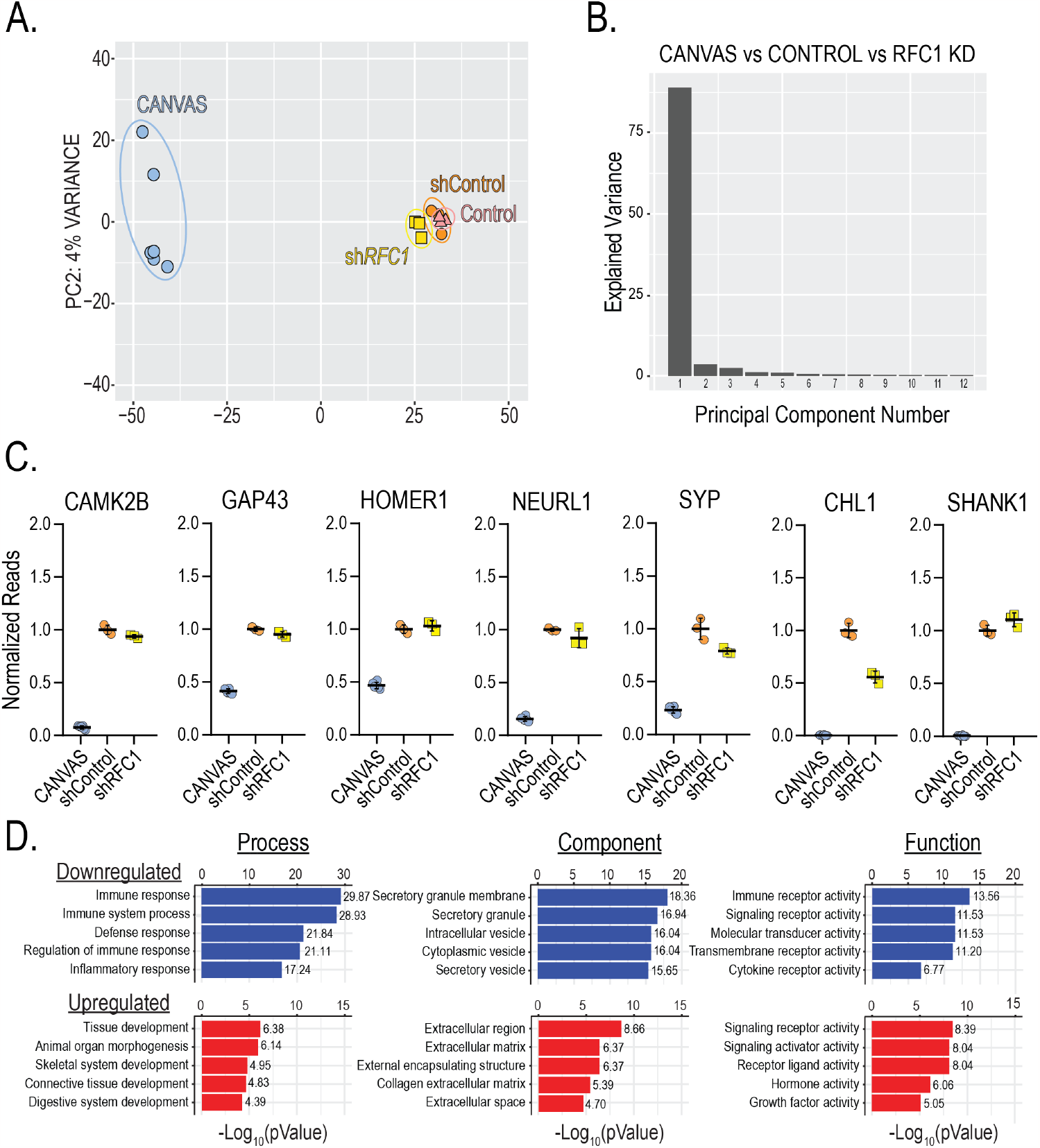
Supplemental data to RFC1 knockdown RNASeq analyses. **A**) Principal component analyses of CANVAS (n=6), control (n=3), control *shControl* (n=3), and control *shRFC1* (n=3) patient iPSC-derived neurons. **B**) Scree plots of explained variance with principal component axis of variance per sample shown for PC1 and PC2. **C**) Normalized read counts for CAMK2B, GAP43, HOMER1, NEURL1, SYP, CHL1, and SHANK1 from CANVAS (n=6), *shControl* (n=3), and *shRFC1* (n=3) iPSC-derived neurons.

**Supplementary Figure 12.**
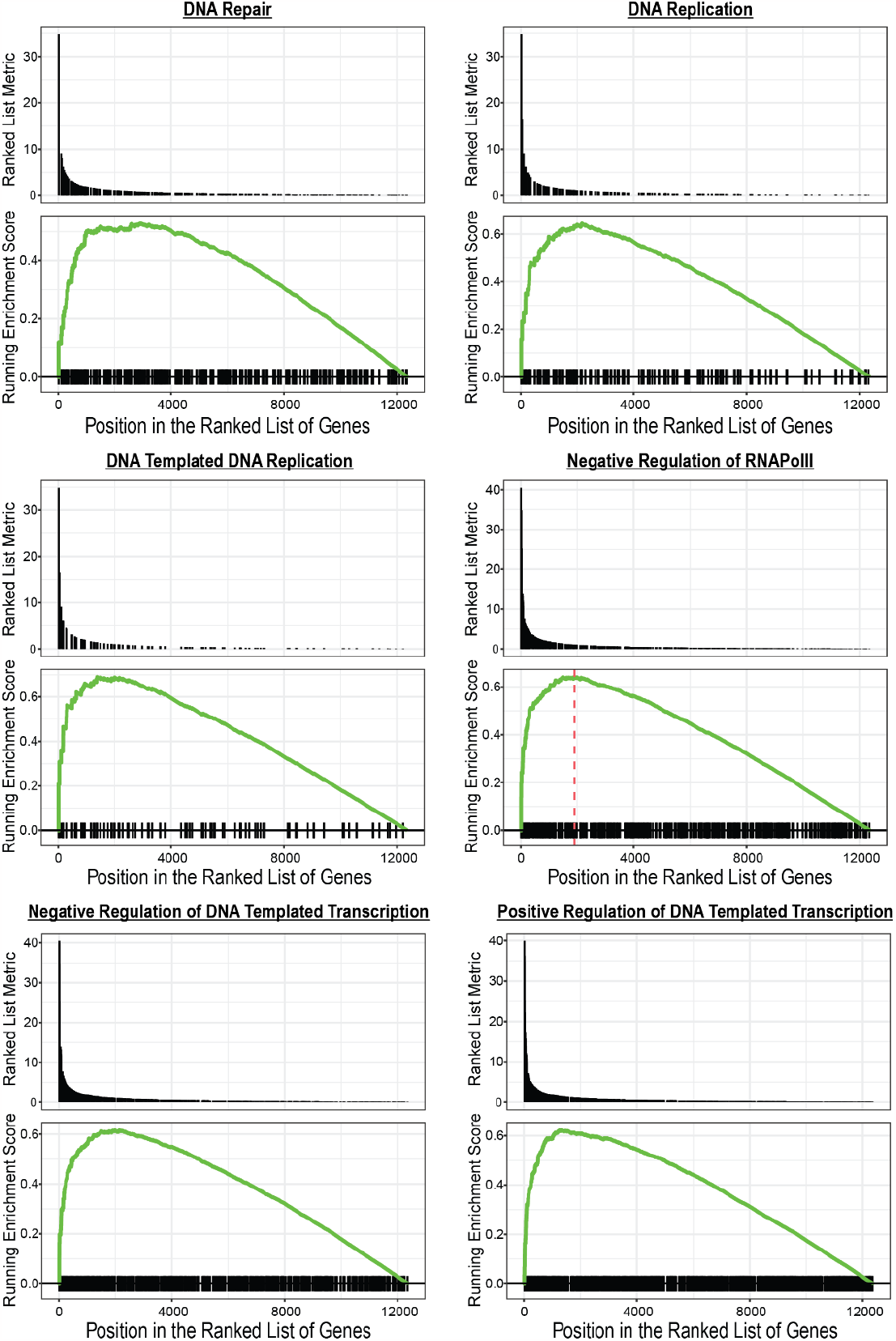
Gene set enrichment analysis (GSEA) of RFC1-associated functions in RFC1 knockdown control iPSC-derived neurons. Gene set enrichment analysis (GSEA) of the RFC1-associated functions DNA repair, DNA replication, DNA templated DNA replication, negative regulation of RNA Pol II, and positive and negative regulation of DNA templated transcription in RFC1 knockdown control iPSC-derived neurons.

**Supplementary Figure 13.**
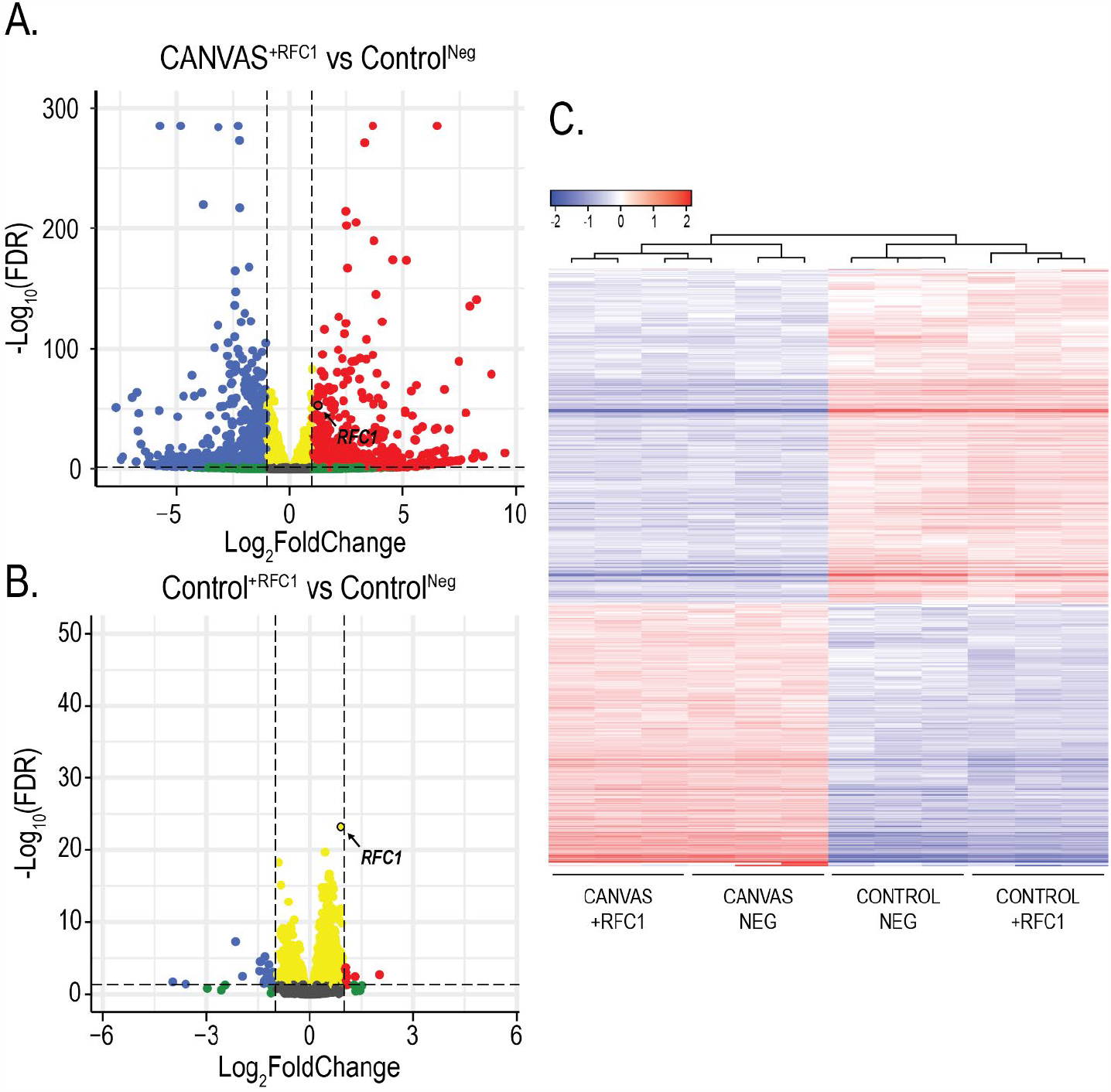
RFC1 reprovision RNASeq analyses. **A**) Volcano plot of -Log_10_FDR vs Log_2_(Fold Change) for CANVAS patient-derived neurons transduced with either full-length RFC1 CDS lentivirus or control lentivirus (n=3/group), RFC1 labelled. **B**) Volcano plot of -Log_10_FDR vs Log_2_(Fold Change) for control-derived neurons transduced with either full-length RFC1 CDS lentivirus or control lentivirus (n=3/group), RFC1 labelled. **C**) Heatmap of normalized expression for the top 1000 genes differentially expressed in CANVAS, control, and CANVAS isogenic patient iPSC-derived neurons.

## Methods

### Dermal fibroblast isolation and iPSC derivation from canvas patient samples

Dermal fibroblasts were obtained from consenting patients clinically diagnosed with CANVAS spectrum disorder and genetically confirmed to possess biallelic *RFC1* expansions. Dermal biopsy samples were cultured in Fibroblast Medium (DMEM, 10% (v/v) FBS, 1% (v/v) 100x NEAA, 1% (v/v) Pen/Strep, 1.5% (v/v) 1M HEPES) at 37 ºC and 5% CO_2_ until fibroblasts emerged from the tissue and were maintained at low passage number and grown to 80-90% confluence prior to episomal reprogramming. On day 0, fibroblasts were detached with Trypsin-EDTA, washed with -/- Phosphate Buffered Saline (PBS), and 3x10^5^ cells were resuspended in 120 μl of R-buffer (Invitrogen) supplemented with 1 μg GFP plasmid, 1μg pCXLE-hUL (Addgene #27080), 1 μg pCXLE-hSK (Addgene #27078), and 1 μg pCXLE-hOCT3/4-shP53 (Addgene #27077) and electroporated using the Neon System (Invitrogen, Condition: 1450 V, 10 ms, 3-pulses). Electroporated fibroblasts were subsequently plated across 3 wells of a Geltrex™ (Thermo Scientific) coated 6-well plate with daily media changes of fresh fibroblast media until day 3. On day 3 cells were changed into a 50/50 ratio of fibroblast media and TeSR™ E-7™ Reprogramming Media (STEMCELL Technologies), and on day 5 were changed into 100% E-7™ Reprogramming Media. Daily media changes of E7™ Reprogramming Media followed until the emergence of iPSC colonies between day 21-28 when iPSC colonies were picked and transferred to Geltrex™ coated 12-well plates containing 1 mL TeSR™-E8™ (STEMCELL Technologies) supplemented with 10 μM ROCK Inhibitor (Y-27632, Cayman Chemical) for 24 h. iPSCs were maintained in TeSR™-E8™ at 37 ºC and 5% CO_2_ and passaged with 0.5 mM EDTA as needed. Multiple iPSC colonies were picked, expanded, and characterized per patient line before use. iPSC colonies were stained via ICC for pluripotency markers using antibodies against SOX2 (ab5603), OCT4 (ab181557) & Nanog (ab21624), and mRNA expression of pluripotency transcription factors was confirmed by RT-PCR (Supplementary Figure 1A-B). iPSCs were confirmed to have no chromosomal abnormalities by G-band karyotyping (Supplementary Figure 1B, WiCell Research Institute).

### Generation of heterozygous isogenic iPSC lines by CRISPR-Cas9

CRISPR gRNAs were designed to remove the expanded AAGGG repeat by non-homologous end joining (NHEJ) utilizing the closest unique PAM sites to the repeat region and obtained from IDT (Figure 1B). iPSCs were detached to a single cell suspension using Accutase™ (ThermoFisher), and 1x10^6^ cells were resuspended in 120 μL R-buffer containing preformed Cas9 RNP complexes of (20 μM HiFi Cas9 (IDT), tracR-ATTO^550^ (IDT), and gRNAs: F: GAGAATAGCAACGGTGTAGCTGG, R: TCATTTTCTGAAATACGGACAGG). The iPSC:RNP mix was electroporated using the Neon system (Condition: 1450 V, 10 ms, 3-pulses), before plating across 3 wells of a Geltrex™ coated 6-well plate with TeSR™-E8™ media supplemented with 10uM ROCK inhibitor (Y-27632) for 24 h. Electroporation efficiency was assessed by nuclear positive fluorescence of TracR ATTO^550^, and iPSCs were cultured until healthy colonies emerged. Clonal populations were achieved through single-cell isolation and expansion^62^. Emerging clonal colonies were screened by PCR utilizing primer sets that spanned the repeat, priming either inside or outside the deletion region (Figure 1C, Supplementary Table 1). Utilizing these primer sets, lack of amplification using primer set 1 in conjunction with amplification using primer set 2 indicates a biallelic expansion at this locus, amplification with both primer sets indicates either a biallelic wild-type locus, or the presence of a heterozygous monoallelic expansion when combined with a positive saw-tooth like pattern by repeat-primed PCR. Finally, reduced molecular weight amplification utilizing primer set 1 and full-length amplification with primer set 2 in combination with a positive saw-tooth like pattern by repeat-primed PCR indicates a monoallelic deletion of this locus by CRISPR-Cas9, and reduced molecular weight amplification with primer set 1 and absence of amplification with primer set 2 indicates a biallelic deletion of this locus byCRISPR-Cas9. Successful deletion of a single allele was achieved in CANVAS patient 1 iPSCs (Figure 1C), and the specificity of this deletion was investigated by sanger sequencing of long-range PCR products (Figure 1D).

### *RFC1* repeat screening and repeat-primed PCR

Cell pellets were obtained from patient-derived cell lines by Accutase™ detachment and centrifugation, or from patient post-mortem brain tissue through gentle neutral protease digestion of brain tissue chunks for 45 mins at 37 °C, followed by mechanical dissociation to single cell slurry by trituration. Genomic DNA was subsequently isolated from cell pellets using gDNA mini-prep kit (Zymo). Screening for repeat expansions in *RFC1* was achieved by a combination of short-range repeat-spanning end-point PCR (Supplementary Table 1) with 2X Faststart PCR mastermix (Roche) and by Repeat-Primed PCR using 2X Phusion Flash High-Fidelity PCR Mastermix (Thermo Fisher). End-point PCR was achieved by amplifying across the repeat region where the presence of a single band at the expected size indicated a non-expanded Wild-Type locus, whereas the absence of a band in the presence of a control band utilizing primers that amplified the intronic region adjacent to the repeat locus indicated the presence of a large, expanded region (Figure 1C). Repeat primed PCR of the repeat locus was conducted as previously described^8^. 5’-FAM labelled PCR products were analyzed through capillary electrophoresis by Laragen Inc. and files were analyzed by Peak Scanner Fragment Analysis Software (Figure 1E). Primers and cycling conditions for all PCR/RP-PCR experiments are outlined in Supplementary Table 1.

### Differentiation and maintenance of patient iPSC-derived neurons

iPSCs were differentiated to neural precursor cells (NPCs) and glutamatergic forebrain neurons using a modified dual-SMAD inhibition protocol^63^ (Supplementary Figure 1C-E). Briefly, iPSCs were detached and plated as a monolayer at 100% confluence in TeSR™-E8™ media containing 10 μM ROCK Inhibitor (Y-27632) for 24 h. On day 2, media was changed to 3N+A Media (241 mL DMEM/F12 with HEPES + L-Glutamine, 241 mL Neurobasal Medium, 5 mL NEAA, 2.5 mL N2 supplement, 5 mL B27 (- Vitamin A), 2.5 mL Glutamax™, 2.5 mL Pen/Strep, 125 μl Insulin, and 3.5 μl BME) supplemented with two SMAD inhibitors: 10 μM SB431542 (Cayman Chemical) and 1 μM dorsomorphin (Cayman Chemical). At day 10-12, neuroepithelial sheets were combed into large clumps, passaged, and maintained on Geltrex™ coated plates in 3N+A media supplemented with 20 ng/mL Fibroblast Growth Factor (FGF), with daily media changes until neuronal rosettes appeared (Supplementary Figure 1D). Rosettes were manually picked and dissociated into single cell NPCs using Accutase™ and plated on to Geltrex™ coated plates in neural expansion medium (3N+A containing 20 ng/mL FGF and 20 ng/mL epidermal growth factor (EGF)) with media changes every other day and passaged as needed using Accutase™. Neuronal differentiation was achieved by plating high-density NPCs and switching to Neuronal Induction Media (490 mL Brainphys, 10 mL SM1, 5 mL N2, 20 ng/mL Brain Derived Neurotrophic Factor (BDNF), 20 ng/mL Glial Derived Neurotrophic Factor (GDNF), 200 nM L-ascorbic acid, and 1 mM bucladesine (dbcAMP)). Neurons were maintained with half medium changes twice per week, with Neuronal Induction Media supplemented with 1 μg/mL laminin once per week. After 3 weeks of neuronal induction, post-mitotic neurons were detached using Accutase™ and replated at a uniform density on PEI & laminin coated plates before experimentation.

### Lentivirus production and transduction of iPSC-derived neurons

pLV-eF1-RFC1-eGFP full-length RFC1 expression plasmid was designed and synthesized by VectorBuilder, and pSMART-eF1-RFC1shRNA-tGFP expression plasmids were designed and produced by Horizon Discovery (Table 1). Plasmids were prepped for lentiviral production by bacterial transformation followed by maxi-prep. 80 μg of plasmid DNA was used for lentiviral particle generation at University of Michigan Viral Vector Core. Lentivirus particles were delivered resuspended in DMEM media at 100x concentration (1x10^8^ TU/mL). Lentivirus transduction of patient iPSC-derived neurons occurred on DIV7 post-differentiation: neurons were first treated with 1 μg/mL polybrene for 30 minutes, before being washed with fresh complete Brainphys media and subsequently changed to conditioned media containing lentiviral particles diluted to their working concentration. Neurons were changed to fresh media 24 h after transduction, and fluorescently positive cells appeared ∼72 h post-transduction. Efficiency of lentiviral transduction is assessed in supplementary figure 7.

### Generation of repeat-containing CANVAS DNA reporter plasmids

Plasmids containing expanded AAGGG/AAAAG or CCCTT/CTTTT repeats were generated using Recursive Directional Ligation (RDL)^29^. Briefly, gene blocks containing 16 repeats flanked by type IIS restriction sites were produced by Genewiz (Azenta), and these were ligated into pcDNA3.1 plasmids mutated to remove restriction sites of interest. The repeat size was recursively doubled through a process of plasmid digestion to remove the repeat unit, followed by re-ligation back into the donor plasmid that had been nicked open at the 3’ end of the repeat unit. When the repeats were built to the maximal length before significant repeat shrinkage was observed, the repeats were removed by restriction endonuclease digestion, and ligated into pcDNA3.1 reporter plasmids containing 100 nt of sense or antisense intronic sequence (Figure 2A).

### Longitudinal survival assays of primary rodent and iPSC-derived neurons

Mixed cortical neurons were dissected from E20 Long–Evans rat pups of both sexes, as previously described^35,36^. Cortical neurons were cultured at 0.6 × 10^6^ cells mL^−1^ on 96-well plates. Cultures were maintained at 37 °C in Neuronal Growth Media (NGM - Neurobasal-A supplemented with 2% (v/v) B-27 and 1% (v/v) Glutamax™ (Fisher)). On DIV 4, neurons were co-transfected with 0.1 μg of pGW1-mApple and either 0.1 μg of pGW1-GFP or 0.1 μg of experimental DNA per well of a 96-well culture plate, using Lipofectamine 2000 (Invitrogen). Neurons were imaged at regular 24 h intervals starting 24 h post-transfection using an automated fluorescence microscopy platform detailed in earlier studies^35,36^. Image processing for each time point and survival analysis for automated fluorescence images were achieved by custom code written in Python or the ImageJ macro language, and cumulative hazard plots were generated using the survival package in R.

### Post-mortem brain IHC and antibody generation

Rabbit polyclonal antibodies were generated by Abclonal (Cambridge, MA) against peptides containing (KGREG)_7_ or (PFPSL)_7_, corresponding to (AAGGG)_21_ and (CCCTT)_21_ respectively. Antibodies were affinity purified from anti-sera. Pre-bleed sera and peptides were used to characterize the antibodies and protein concentrations of the pre-bleed sera were determined. The dilution of pre-bleed sera used had an equal protein concentration as the dilution of antibody used. A 100x excess of peptide was incubated with the antibody prior to use to block the antibodies’ ability to bind to antigen.

CANVAS and control brain regions were obtained from the University of Michigan Brain Bank, Harvard University Brain Bank, and the University of Amsterdam Brain Bank with informed consent of the patients or their relatives and the approval of the local institutional review boards (IRB). For immunohistochemistry, paraffin embedded sections were de-paraffinized with a series of xylene washes and decreasing concentrations of ethanol. Antigen retrieval (AR) was achieved with citrate buffer, if necessary. Endogenous peroxidase was quenched with 1% hydrogen peroxide for 30 min. Sections were blocked in 5% normal goat serum (NGS) in Tris, pH 7.6. Primary Antibodies, KGREG_7_ (Abclonal, 1:100, Acid AR), PFPSL_7_ (Abclonal, 1:100, Acid AR) were diluted in 5% NGS Tris B (Tris pH 7.6, 0.1% Triton-X 100, 0.5% bovine serum albumin) and were incubated with sections overnight at 4 °C. The following day, sections were washed in Tris A (Tris pH 7.6, 0.1% Triton) and Tris B. Antibody detection was determined using VECTASTAIN Elite ABC HRP kit (Vector Laboratories) following the manufacturer’s protocol. Sections were counterstained with hematoxylin (Vector Laboratories). After dehydrating the slides through a series of increasing concentration of ethanol and then xylenes, coverslips were mounted to the slides using DPX mounting media. Images were taken on an Olympus BX51 microscope.

### RNA hybridization chain reaction (HCR) of RNA foci

CANVAS patient and control iPSC-derived neurons were differentiated and maintained to 8 weeks of age as previously described. Cells were washed with (-/-) PBS and subsequently fixed with 4% paraformaldehyde (PFA) and incubated overnight in 70% ethanol. After overnight incubation with 70% ethanol, cells were rehydrated in PBS for 1 h, permeabilized with 0.1% Triton X-100 for 5 min, and blocked with 2% bovine serum albumin (BSA) for 20 min at room temperature. Repeat-containing RNA foci were probed in patient iPSC-derived neurons using DNA probes with additional sequence complementary to Cy5-labeled, self-hybridizing hairpins^64–67^. Probes against the AAAAG, AAGGG, CTTTT & CCCTT sequences (Supplementary Table 2) were purchased from molecularinstruments.org and applied according to the manufacturer’s protocol. Coverslips were then applied to slides with ProLong Gold Antifade Mounting Medium with DAPI. Then, 10– 20 fields per condition were imaged blinded using an inverted, Olympus FV1000, laser-scanning confocal microscope. Channels were imaged sequentially and optimized to eliminate bleed-through. Neurons were imaged in a series of Z-planes to resolve the entire soma and dendritic arbor. Images were analyzed blinded in ImageJ. Average intensity composite images were derived from raw image files. For quantification of individual soma, cell casts were made using threshold images as a guide.

### Analysis of DNA damage accumulation and recovery in iPSC-derived neurons

The accumulation of basal DNA damage was analyzed in 8-week-old patient iPSC-derived neurons by Western Blot for the DNA damage marker γ-H2AX (P-S139, Abcam ab11174). The ability to detect DNA damage was tested in patient iPSC-derived neurons by exposure to 0, 15, 30, and 120 mJ/cm^2^ ultra-violet (UV) irradiation to induce differing degrees of DNA damage, followed by Western Blot analysis for γ-H2AX. Recovery after DNA damage induction in CANVAS patient iPSC-derived neurons was conducted as described: CANVAS patient and control iPSC-derived neurons were plated on PEI/laminin coated glass chamber slides and maintained for 8 weeks as previously described. At 8 weeks, neurons were either fixed with 4% PFA for no-exposure control or irradiated with 60 mJ/cm^2^ UV. UV Irradiated cells were fixed in 4% PFA at time intervals of: 0.5, 1, 3, 6, 12, and 24 h post-irradiation, and were stained for the neuronal nuclei marker NeuN, DNA marker DAPI and DNA damage marker γ-H2AX. Images were captured in all 3 fluorescent channels at the same exposure for each cell line and time point at 40x magnification with an Olympus IX71 fluorescent microscope and Slidebook 5.5 software, and γ-H2AX reactivity analyzed using FIJI. High throughput quantification of cells was achieved by using a FIJI script to mask NeuN positive nuclei using the parameters of size: 200-1500 pixels^2^ and circularity: 0.40-1.00. Masked NeuN positive nuclei were subsequently analyzed for γ-H2AX intensity and was quantified and visualized using GraphPad Prism v.9.

### Cell proliferation assays

CANVAS patient and control fibroblasts were analyzed for cell proliferation rates through automated longitudinal brightfield microscopy using the Incucyte S3 system. Fibroblasts were seeded into TPP™ 96-well plates at a density of 2,500 cells per well and whole wells were imaged by tiling of 5x images per well using S3 Neuro O/NIR Optical Module 10x magnification with 6 h intervals for a period of 10 days. Cell confluency was automatically analyzed using the Incucyte^®^ S3 Live Cell Analysis Software using the basic analyzer cell confluence settings to mask and quantify individual cell number and absolute confluence with the parameters: segmentation adjustment – 2, hole fill – 0 μm^2^, adjust size – 4 pixels. 3 control and 4 CANVAS patient fibroblast lines were used, with 12 wells averaged per fibroblast line for technical replicates within each biological replicate, and a total of 3 biological replicates presented. For lentiviral experiments, fibroblasts were transduced with lentiviral particles in 10 cm dish mass culture 7 days prior to re-seeding at experimental density in 96-well plates. Lentiviral particles were diluted within culture medium to a final concentration of 1x.

### Calcium imaging and analysis of iPSC-derived neurons

TPP™ tissue-culture treated 96-well plates were prepared by coating with 0.1% polyethyleneimine (PEI) diluted in borate buffer for 24 h, followed by 3x washes in H_2_O and coating with 1 μg/mL laminin diluted into neuronal medium overnight before use. Rat primary cortical astrocytes (Fisher) were plated at high density and maintained until confluence in Astrocyte Media (DMEM/F12, 10% (v/v) Horse Serum, 10% (v/v) FBS, 3% (v/v) 20% D-glucose solution, 2% (v/v) B27 supplement, 1% (v/v) Glutamax™, 1% (v/v) Pen/Strep antibiotic). CANVAS and control iPSC-derived neurons were differentiated from NPCs in Geltrex™-coated 10 cm plates as previously described for 21 days. 5 days post-differentiation, neurons were transduced with lentiviral particles as previously described based on experimental setup. After 3-weeks neuronal differentiation, neurons were detached using 25% Accutase™ and incubating at 37 ºC for 10 mins. Neurons were diluted in pre-warmed Brainphys neuronal media and gently triturated to detach. Neurons were pelleted by centrifugation and resuspended in 1 mL pre-warmed Brainphys neuronal media before passing through a 45 μm cell strainer to obtain a single cell suspension of neuronal soma. Cells were counted using a hemocytometer with trypan blue as a live cell marker, and neurons were diluted to a concentration of 5,000 live cells in 200 μl complete neuronal media per well of a 96-well plate.

10 days post-replating, neurons were transduced with Incucyte^®^ Neuroburst Orange Lentivirus particles by diluting 5 μl lentivirus into complete Brainphys neuronal media per well, and neurons were maintained as previously described until analysis using the Incucyte^®^ S3 Live Cell Analysis Instrument. Image acquisition was conducted using S3 Neuro O/NIR Optical Module at 4x magnification for 180 s per well in phase and orange fluorescence channels, and data were automatically analyzed using the Incucyte^®^ S3 Live Cell Analysis Software with the parameters: object size - 30 μm, min cell width - 9 μm, sensitivity - 1, min burst intensity - 0.2. Neurons were analyzed for active cell number, basal calcium intensity, burst rate, burst strength, durst duration, and network correlation. Data for different neuronal signaling metrics were compiled for each scan time point and analyzed using GraphPad Prism v.9.

### RNA extraction and RT-PCR/qPCR

Total RNA was extracted using TRIzol and chloroform, followed by isopropanol precipitation. RNA pellets were washed in 75%+ ethanol and resuspended in nuclease free H_2_O. Contaminating gDNA was removed by DNase I digestion (Fisher Scientific) and cleaned up using an RNA clean & concentrator kit (Zymo). Total RNA was quantified by NanoDrop UV/Vis spectrophotometry, and 1 μg of RNA was reverse-transcribed (RT) to cDNA using iScript cDNA synthesis kit (Bio-Rad) with oligo-dT primers for poly-A mRNA selection. RT-PCR was achieved using 2X Faststart PCR mastermix (Roche) using primer sets and cycling conditions described in Supplementary Table 1. RT-qPCR was achieved using Taqman Real-Time PCR Assay(s) (Thermo Fisher) using primer probe sets and cycling conditions described in Supplementary Table 1.

### RNA-seq sample preparation and analysis

For RNA-seq experiments, patient iPSC-derived neurons were plated and aged to ∼10 weeks as previously described. Total RNA was extracted using TRIzol and chloroform, followed by isopropanol precipitation. RNA pellets were washed in 75%+ ethanol and resuspended in nuclease free H_2_O. Contaminating gDNA was removed by DNase I digestion (Fisher Scientific) and cleaned up using an RNA clean & concentrator kit (Zymo). Total RNA was quantified by NanoDrop UV/Vis spectrophotometry, analyzed for integrity by Bioanalyzer, and 1.5-2 μg total RNA was sent to Genewiz (Azenta) for rRNA-depletion library preparation and paired-end (PE) sequencing, where between 20-30 M paired-end reads were achieved per sample. Raw FASTA read files were delivered by sFTP transfer and analyzed in Command Line and R-Studio. Read files were quality controlled using *FastQC(v0*.*12*.*0)*^68^, pre-processed and trimmed using *fastp(v0*.*23*.*4)*^69^, and aligned to GRCh38 (NCBI GCF_000001405.40) genome using *STAR(v2*.*7*.*11a)* two-pass alignment^70^. Transcript and gene count matrices were produced using *featureCounts(Subread v2*.*0*.*6)*^71^. Read count matrices were imported into R-Studio (vRtidyverse/4.2.0) where count normalization and differential gene expression analysis was conducted using *DESeq2(v1*.*40*.*2)*^72^. Gene Ontology (GO) Pathway Overrepresentation analysis was conducted using *GOstats(v2*.*66*.*0)*^73^, and Gene Set Enrichment Analysis (GSEA) conducted using *clusterProfiler(v4*.*8*.*3)*^74^ and *gseGO(v3*.*0*.*4)*^74^. Plots were generated and visualized using *DESeq2(v1*.*40*.*2)*^23^, *EnhancedVolcano(v1*.*18*.*0)*^75^, *ggPlot2(v3*.*4*.*3)*^76^, and *ggRidges(v0*.*5*.*4)*^76^. Differential exon usage was analyzed and plotted using *DEXSeq(v1*.*46*.*0)*^51^, circular RNA analysis was conducted using *DCC(v0*.*5*.*0)* in Python^77^. Sashimi plots of splice junction analysis were produced using Integrative Genomics Viewer (IGV) Genome Browser^78^ and exported for modification in Illustrator (Adobe). All harvesting and sequencing of RNA was conducted at the same time within each experimental condition to avoid batch effects. Genes and pathways that showed significant correction in the heterozygous isogenic corrected line from CANVAS to control were identified by binning normalized gene expression values into categories of those showing reduced expression upon heterozygous deletion, 0-50% positive correction in expression compared to control, 50-150% positive correction in expression compared to control, and those genes that showed over 150% correction in comparison to control. These genes were further binned into their statistical significance in difference of expression when compared to control and plotted using *ggPlot2(v3*.*4*.*3)* ^*ref*^ and Gene Ontology (GO) Pathway Overrepresentation analysis was conducted using *GOstats(v2*.*66*.*0)*^*ref*^.

## Author Contributions

Conceptualization, **C.J.M**. and **P.K.T**.; methodology, **C.J.M**., **A.K**., **S.J.G**., **Y.P**., **J.M**., and **A.S**.; investigation, **C.J.M**., **A.K**., **Y.P**., **J.M**., **A.S**., **M.A**., and **A.A.D**.; formal analysis, **C.J.M**., **A.K**., and **S.J.G**.; software, **S.J.B**.; resources, **C.J.M**., **V.K**., **A.A.D**., **S.J.B**., and **P.K.T**.; funding acquisition, **P.K.T**. and **S.J.B**.; supervision, **P.K.T**.; visualization, **C.J.M**.; writing – original draft, **C.J.M**. and **P.K.T**.; writing – review & editing, **C.J.M**., **A.K**., **S.J.G**., **Y.P**., **J.M**., **A.S**., **M.A**., **V.K**., **S.J.B**., **A.A.D**., and **P.K.T**.

## Acknowledgements

This work was supported by funding from the National Institute of Neurological Disorders and Stroke (R21NS129096-01, R01NS086810-09 to **P.K.T**; R01NS097542, R01NS113943, and R56NS128110 to **S.J.B**), National Institute of Child Health and Human Development (P50HD104463-04). **P.K.T** was also supported by Veterans Health Administration (BLRD BX004842) and philanthropic donations. **Y.P** and **A.S** were supported by NIH Postbaccalaureate Research Education Program (PREP, PAR-22-220 R25). **J.M** was supported by the UM Medical Scientist Training Program (MSTP). We wish to thank the patients and their families for the donation of fibroblast and post-mortem tissue samples for this study. We also thank the University of Michigan Human Stem Cell and Genome Editing Core, Dr Jack Parent, and Sandra Mojica-Perez for their continuous support in generating CRISPR-corrected iPSC clones and for the use of their Incucyte S3 instrument.

## Data and materials availability

RNASeq data will be publicly available upon publication of this manuscript. Code for analyses and any materials used for this study are available upon request.

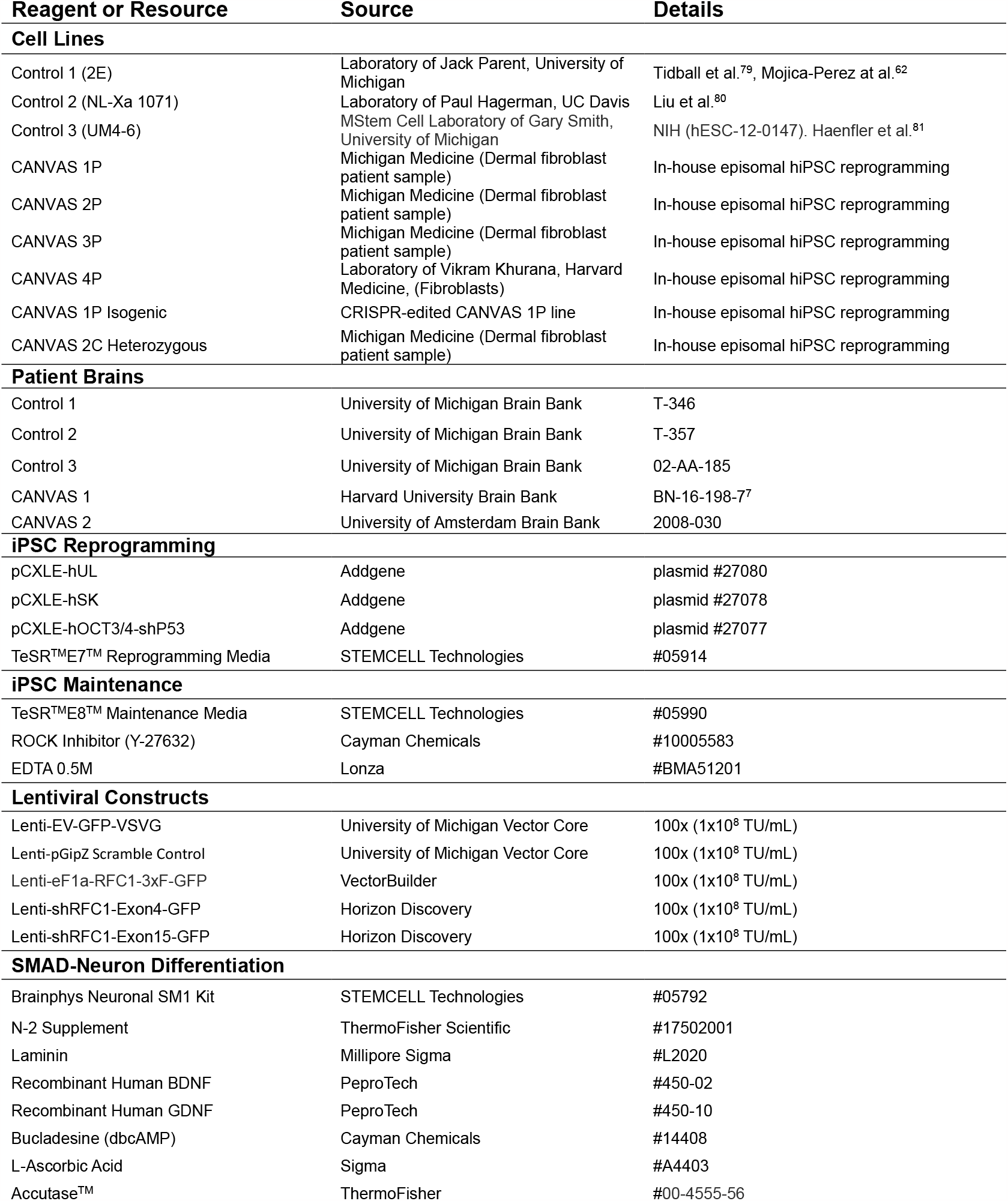
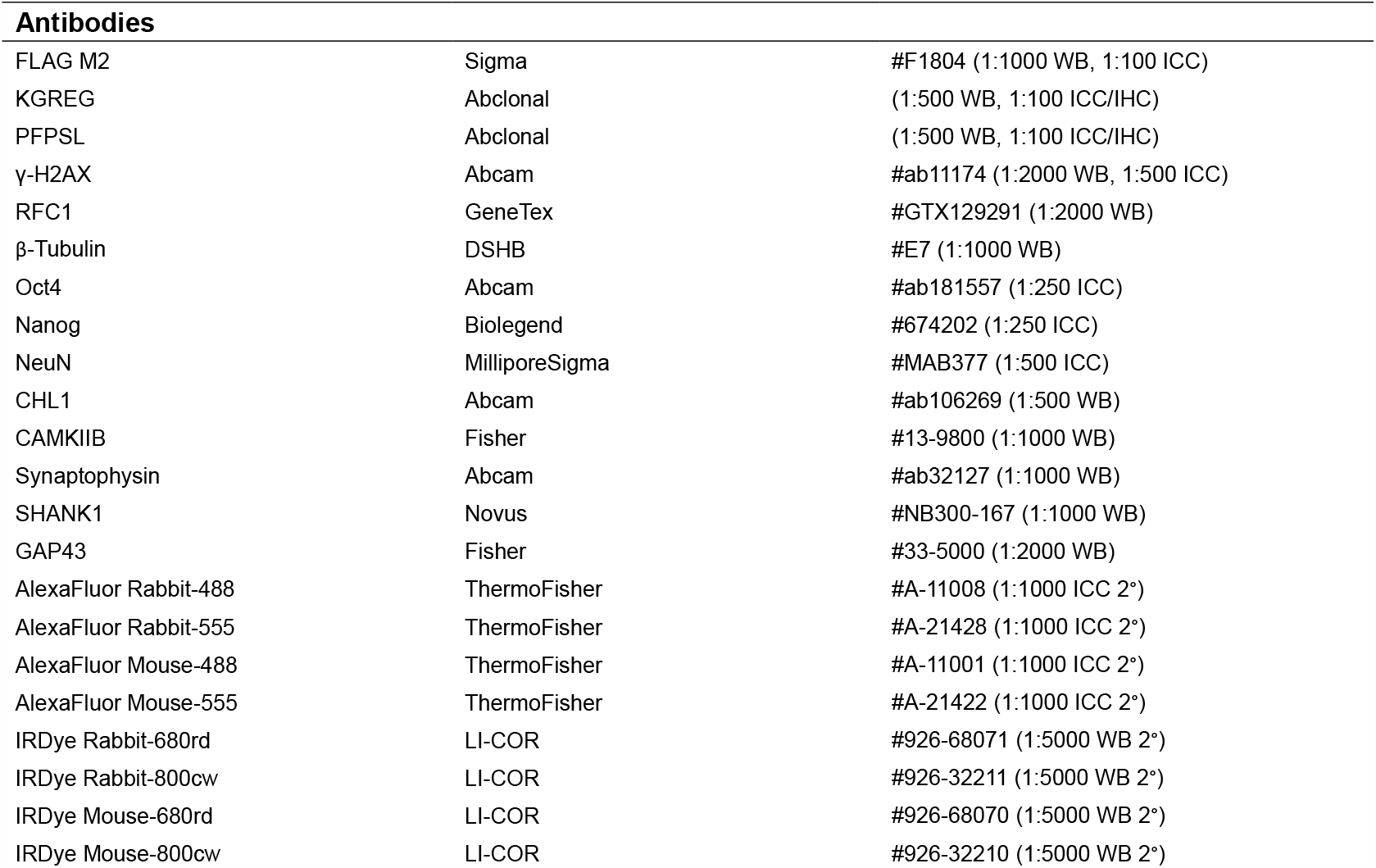

**Supplementary Table 1.**
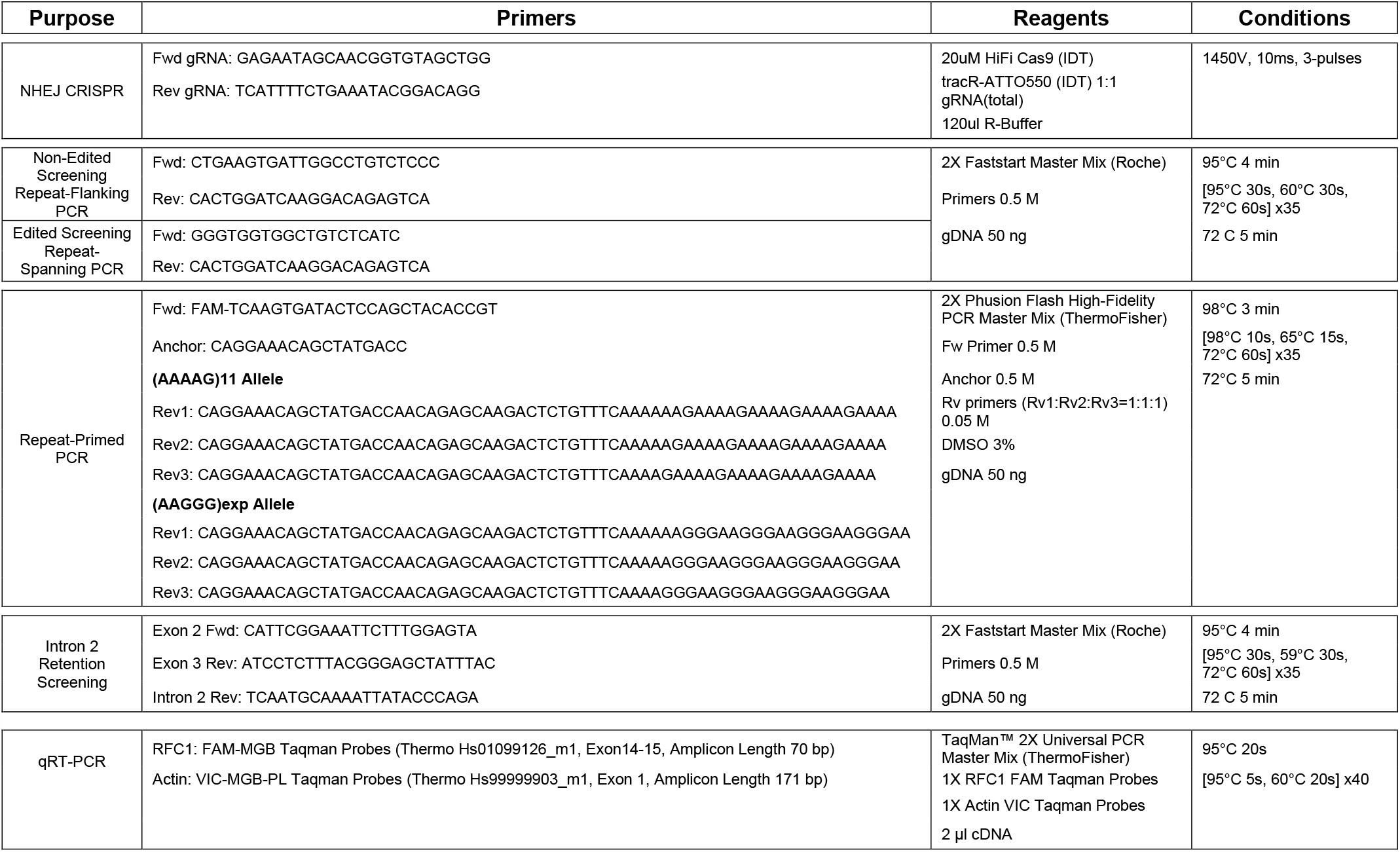
Table of PCR primer sequences, reagents, and thermocycling conditions used.

**Supplementary Table 2.**
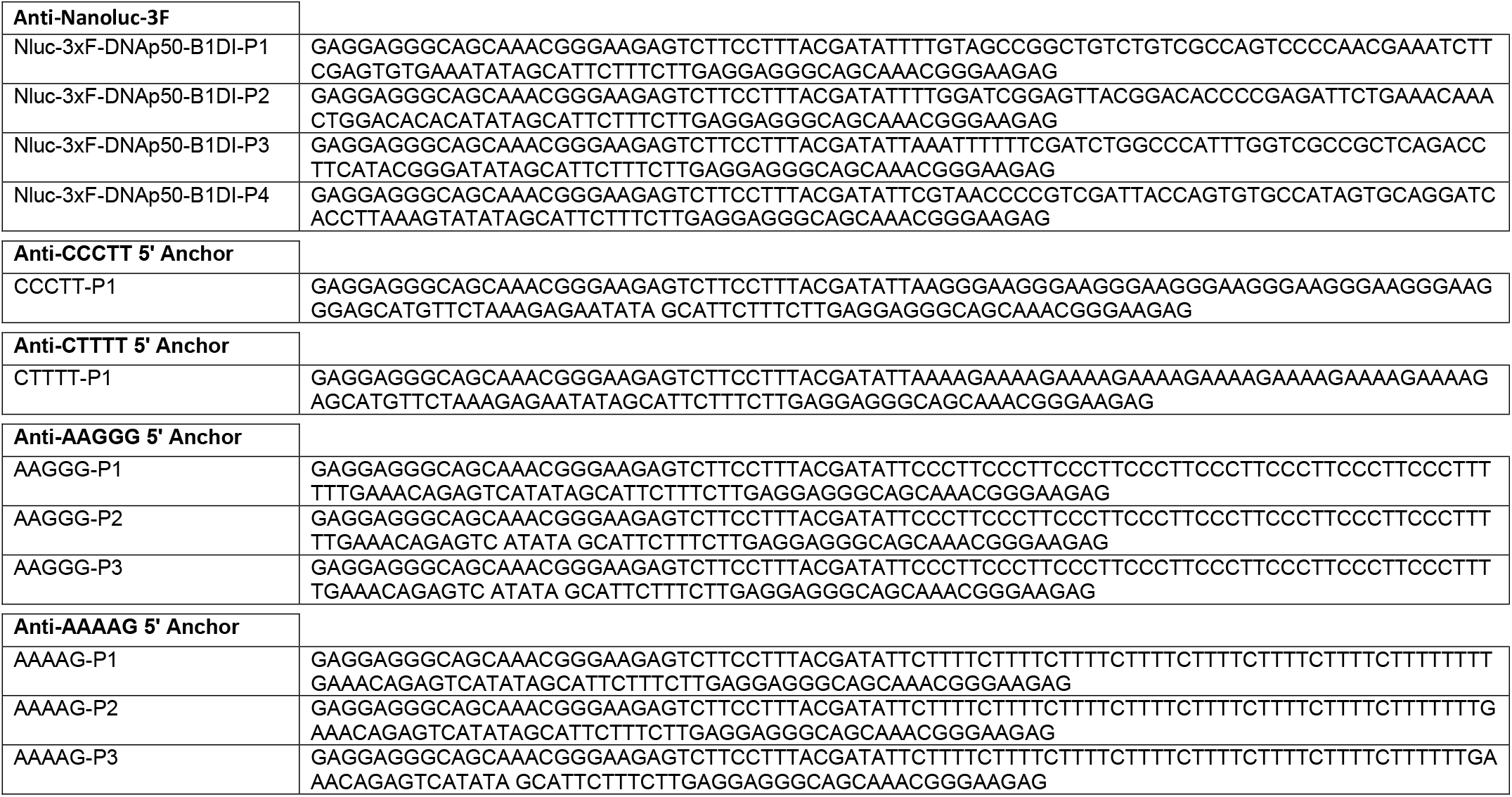
Table HCR probe sequences used.

